# Influence of Sequencing Technology on Pangenome-level Analysis and Detection of Antimicrobial Resistance Genes in ESKAPE Pathogens

**DOI:** 10.1101/2025.01.08.631980

**Authors:** Alba Frias-De-Diego, Manuel Jara, Cristina Lanzas

## Abstract

As sequencing costs decrease, short-read and long-read technologies are indispensable tools for uncovering the genetic drivers behind bacterial pathogen resistance. This study explores the differences between the use of short-read (Illumina) and long-read (Oxford Nanopore Technologies, ONT) sequencing in detecting antimicrobial resistance (AMR) genes in ESKAPE pathogens (*Enterococcus faecium, Staphylococcus aureus, Klebsiella pneumoniae, Acinetobacter baumannii, Pseudomonas aeruginosa,* and *Enterobacter cloacae*). Utilizing a dataset of 1,385 whole genome sequences and applying commonly used bioinformatic methods in bacterial genomics, we assessed the differences in genomic completeness, pangenome structure, and AMR gene and point mutation identification. Illumina presented higher genome completeness, while ONT identified a broader pangenome. Hybrid assembly outperformed both Illumina and ONT at identifying key AMR genetic determinants, presented results closer to Illumina’s completeness, and revealed ONT-like pangenomic content. Notably, Illumina consistently detected more AMR-related point mutations than its counterparts. This highlights the importance of method selection based on research goals. Differences were also observed for specific gene classes and bacterial species, underscoring the need for a nuanced understanding of technology limitations. Overall, this study reveals the strengths and limitations of each approach, advocating for the use of Illumina for common AMR analysis; ONT for studying complex genomes and novel species, and hybrid assembly for a more comprehensive characterization, leveraging the benefits of both technologies.

**Impact Statement:** This study provides a comprehensive comparison of short-read (Illumina) and long-read (Oxford Nanopore Technologies, ONT) sequencing technologies in the context of antimicrobial resistance (AMR) detection in ESKAPE pathogens. By analyzing a large dataset of 1,385 whole genome sequences, the research offers valuable insights into the strengths and limitations of each approach, as well as the benefits of the novel approach of hybrid assembly. These findings have broad utility across microbiology, genomics, and infectious disease research. In particular, they apply to the work of researchers and clinicians dealing with AMR surveillance, investigation into outbreaks, and bacterial genome analysis. Given the nuance with which technological differences in genomic completeness, pangenome structure, and AMR determinant detection have been explored in this study, it is a good basis for informed method selection for future research. While the output represents an incremental advance, its significance lies in its practical implications. It thus enables researchers to take more reasonable decisions in designing genomic studies of bacterial pathogens by showing the complementarity of various sequencing approaches and their specific strengths. This could lead to more accurate and comprehensive detection of AMR, which would contribute ultimately to improved antibiotic stewardship and public health strategies.

**Data Summary:** The authors confirm all supporting data, code and protocols have been provided within the article or through supplementary data files.

**Repositories:** All the sequences used for this study are publicly accessible from GenBank, and their individual IDs are disclosed in Supplementary Table 1.

## Background

An estimated 4.95 million annual deaths are caused by antimicrobial-resistant pathogens worldwide ^1^. These deaths are commonly caused by a group of pathogens identified as ESKAPE for their ability to resist antibiotics and “escape” treatment, creating an ongoing problem for modern medicine ^2^. This group of bacteria includes *Enterococcus faecium*, *Staphylococcus aureus, Klebsiella pneumoniae, Acinetobacter baumannii, Pseudomonas aeruginosa,* and *Enterobacter* species ^2, 3^. ESKAPE pathogens are known to cause healthcare-associated infections (HAIs) ^4^. The compromised health statuses of patients in healthcare settings make them particularly vulnerable to infections caused by these pathogens ^5^. However, ESKAPE pathogens can also be transmitted within community settings, making them widely distributed throughout populations ^6^.

Next-generation sequencing technologies have emerged as a critical component of clinical microbiology and antimicrobial resistance surveillance. Whole genome sequencing (WGS) of clinical isolates is extensively used for pathogen typing, resistance gene identification, and evolutionary and disease transmission studies ^7–12^. At the heart of this technological revolution lie two types of sequencing technology with distinct strengths and weaknesses: short-read and long-read sequencing. Short-read sequencing, typified by platforms like Illumina, furnishes rapid and cost-effective data acquisition ^13, 14^. This technology achieves high-depth coverage, or the average number of times each nucleotide or region of interest is sequenced in a genome, facilitating the identification of prevalent genetic variants and mutations within populations. However, its limited read length can pose challenges in accurately reconstructing complex genomic regions, such as repetitive elements, which are integral to understanding pathogen evolution ^13, 15^. In contrast, long-read sequencing, epitomized by rapidly growing platforms such as Oxford Nanopore Technologies (ONT), generates extensive reads that span challenging genomic segments ^16, 17^. This characteristic empowers the accurate delineation of repetitive regions, mobile genetic elements, and structural variations that are instrumental in shaping the genomic landscape of pathogens. Moreover, long-read sequencing offers the potential to uncover novel genetic elements and achieve more comprehensive genome assembly ^17–20^. However, long-read sequencing exhibits a comparatively higher error rate in raw sequences than standard Next-Generation Sequencers (NGS) like Illumina ^21–23^. This discrepancy stems from ONT’s challenges in distinguishing between purines (A and G) and pyrimidines (C and T), leading to increased substitution errors, particularly in comparison to transversions ^21, 24^. Additionally, low-complexity regions, homopolymers, and short tandem repeats pose further challenges for ONT, potentially inflating error rates. With a mean global error rate on raw reads estimated at approximately 6%, the accuracy of ONT heavily relies on the performance of its base-calling software ^21^.

An additional impacting factor on both Illumina and ONT assemblies and gene calling is based on the differences in read lengths. In the case of Illumina, shorter read lengths lead to more fragmented assemblies and, thus, less accurate gene calls. Similarly, there are differences between the library preparation and sequencing kits used for ONT, where ligation presents longer fragments than the rapid kits at the tradeoff of missing small plasmids, while newer kits have increased accuracy as compared to older materials ^25–27^.

The potential differences on the use of both Illumina and ONT are already being considered for research on these bacteria, with Illumina more commonly used for molecular characterization, metagenomics, and epidemiological surveillance ^28–30^, and ONT mostly used for real-time detection, assembly of bacterial whole genomes and rapid diagnosis ^31, 32^. However, oftentimes the decision of whether to use short or long-read sequencing is hard to determine since only few articles have quantitatively compared these differences ^33–36^.

Despite the higher error rates of long-read sequencing, the combination of both long and short reads has proven efficient in overcoming this problem and improving the accuracy of the obtained results in a novel method known as hybrid assembly correction, which is increasingly being used for genomic studies ^37, 38^. However, as technology advances, specifically designed bioinformatic pipelines for handling both short and long reads will gradually bridge the gap between the two methods. This study seeks to uncover potential disparities between short-read (Illumina) and long-read (ONT) technologies in detecting and identifying AMR genes on ESKAPE pathogens. In addition, we evaluated the ability of both technologies to describe the pangenome, the complete set of genes within each species composed of both a central or core genome (general for all strains) and an accessory genome (particular of specific strains or even unique) ^39–41^. The accessory genome is known to have a strong influence on phenotypic diversity, including genes related to drug resistance ^42, 43^. For that reason, pangenome analyses are a powerful approach to identify genetic and functional diversity of bacterial populations ^39–41, 44, 45^. The results of this study aim to provide quantitative information that could help researchers decide which approach would be more suitable for their specific research. Here, we collected whole genome sequences of 1,385 isolates from ESKAPE bacteria sequenced by both short and long-read technologies and performed a series of quality control and commonly used genomic analyses to assess the potential differences associated with each specific method.

## Methods

### Data collection and curation

In this study, we collated a comprehensive dataset composed of publicly available Whole Genome Sequences (WGS) pertaining to ESKAPE pathogens (*Enterococcus faecium* (genome size between 2.5 and 3.2 Mb)*, Staphylococcus aureus* (genome size around 2.8 Mb)*, Klebsiella pneumoniae* (genome size between 5.3 and 5.7 Mb)*, Acinetobacter baumannii* (genome size between 3.6 and 4.2 Mb)*, Pseudomonas aeruginosa* (genome size about 6.3 Mb), and *Enterobacter* species (genome size between 4.5 and 5.52 Mb)), generated by the collection of sequences from BioProjects that contained sequencing data for both Illumina and Oxford Nanopore Technologies (ONT). This dataset, comprising a total of 1,385 sequences (after QC, Supplementary Table 1) per sequencer, was acquired from the National Center for Biotechnology Information (NCBI) Sequence Read Archive (SRA) repository ^46^. Isolates were only used for further analyses if they were sequenced by both Illumina and ONT sequencers. Metadata associated with each isolate included NCBI biosample, bioproject ID, sequencer and sequencing depth is available in Supplementary Tables 1 and 2.

### Genome completeness analysis

Genome completeness was assessed using BUSCO (Benchmarking Universal Single-Copy Orthologs) v4.0 ^47^ and comparing the selected sequences after the QC analysis and against the bacteria_odb10 database.

### Bactopia pipeline

A graphical visualization of the complete pipeline is found in Figure 1.

**Figure 1.**
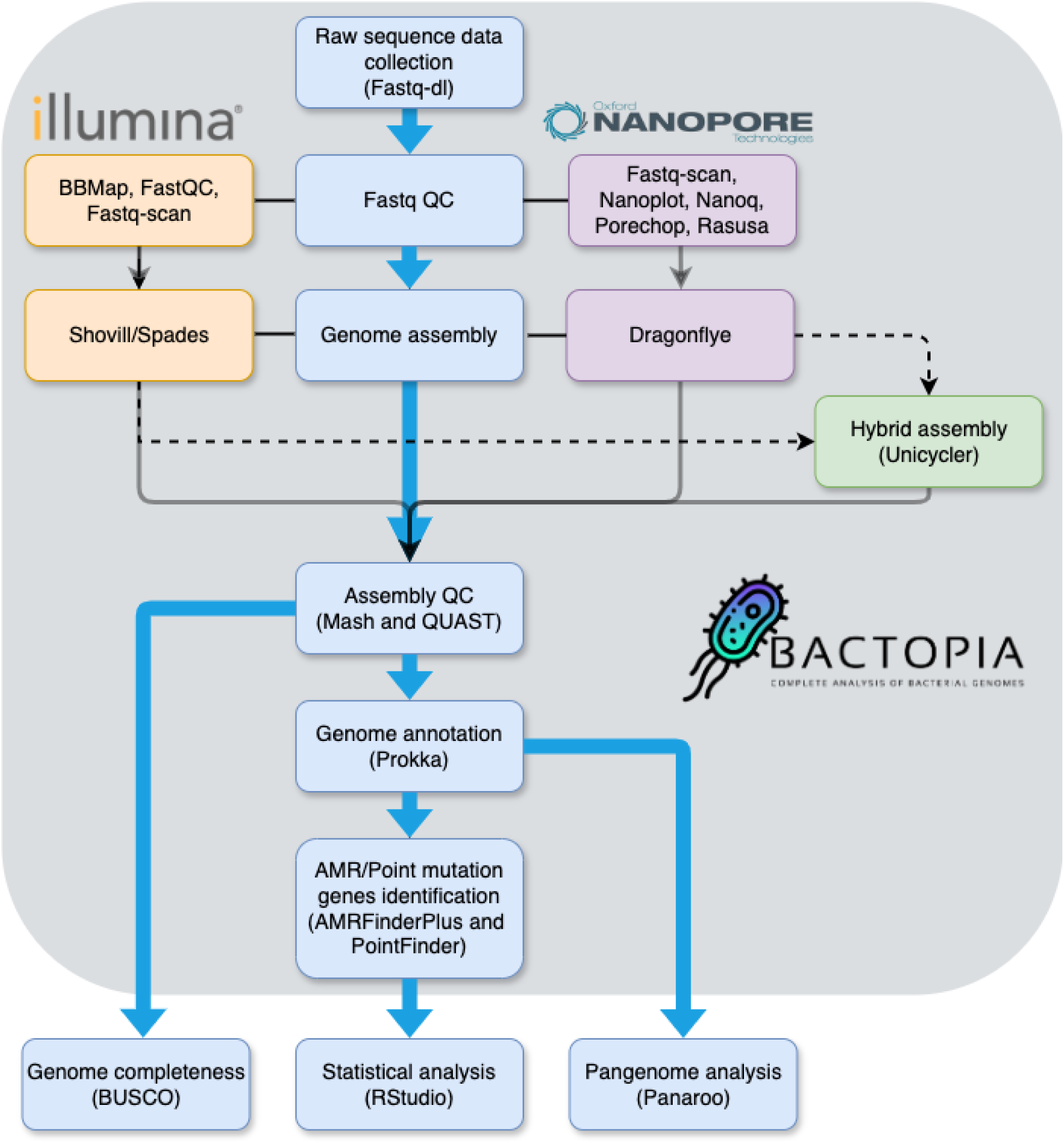
Complete workflow for performing AMR genes identification in ESKAPE pathogens. using a pangenome approach, as well as statistical analysis to identify the potential differences associated with the sequencing technology (e.g., Illumina and Oxford Nanopore Technologies) and hybrid method.

To prepare the raw sequencing data for downstream analysis, we employed the fastq-dl tool v2.0.4 (available fromhttps://github.com/rpetit3/fastq-dl). The raw reads were processed using the Bactopia v3 pipeline ^48^ which uses different tools depending on whether the sequences are short-reads (Illumina) or long-reads (ONT). To ensure high-quality sequencing data, we employed a comprehensive quality control pipeline for Illumina and ONT samples. For Illumina sequences, FastQC v0.12.1 ^49^ provided detailed quality metrics, while BBMap v39.03 ^50^ filtered and trimmed reads with a quality score below Q20. Fastq-scan v1.0.1 (available from https://github.com/rpetit3/fastq-scan) was used for a quick quality overview. For the ONT isolates, Fastq-scan ^51^ provided initial quality summaries, NanoPlot v1.42.0 ^52^ generated detailed quality plots, and Nanoq v0.10.0 ^53^ filtered low-quality reads. Porechop v0.2.4 ^54^ trimmed adapters and split chimeric reads, and Rasusa v0.7.1 ^55^ subsampled reads to a specified coverage. All high-quality reads were then assembled, in the case of Illumina samples using Shovill v1.1.0 ^56^, an integral component of the Bactopia pipeline, which leverages the SPAdes v3.15.5 assembler ^57^, and using Dragonflye v1.1.2 (available from https://github.com/rpetit3/dragonflye) for the ONT samples. To evaluate the assembly quality, assessments were conducted employing Mash v2.3.0 ^58^ and QUAST v5.2.0 ^59^. The genome annotation process was executed using Prokka v1.14.0 ^60^. To leverage the strengths of both short-read and long-read technologies while minimizing their limitations, we performed a hybrid assembly (short-read polishing) using Unicycler v0.5.0 ^61^ in Bactopia. All the annotated sequences (.gff files) were analyzed for AMR gene content using AMRFinderPlus ^62^ as part of the Bactopia pipeline using both nucleotide (BLASTX) and protein (BLASTP) strategies, all matches with identity and coverage below 90% were excluded for further analysis. In addition, point mutations associated with antimicrobial resistance were identified using PointFinder ^63^.

The presence of AMR genes was compared across sequencing methods (Illumina, ONT, Hybrid) using Pearson’s chi-square tests with Yates’ continuity correction implemented in R software ^64^. In addition, the relationship between the presence of AMR genes and point mutations, and the log-transformed total number of bases sequenced per isolate, was assessed through linear regression analysis in R software ^64^.

### Pangenome analysis

To provide an additional level of genomic comparison between the Illumina and ONT isolates, we reconstructed the pangenome of each ESKAPE species using Panaroo (v1.2.2) with the strict clean-mode setting ^65^. To ensure a robust and accurate gene clustering, we set the sequence identity threshold to 95%, meaning that genes needed to have at least 95% sequence similarity to be considered the same gene. This approach allowed us to identify genes that are highly conserved within the population (core genes) as well as those that are more variable (soft-core, shell, and cloud genes). Our threshold for the core genome was set at (>99% of gene similarity among all sequences) to ensure the inclusion of unique gene alleles in the accessory genome. Additionally, we identified the rest of gene types with the following thresholds: soft-core genes (95% - 99%), shell genes (15% - 95%), and cloud genes (<15%).

To further investigate the differences in pangenome size and composition, and assess whether the variations observed were due to differences in annotation, we performed one-way ANOVA analyses to compare the number of annotated genes prior to pangenome reconstruction using R^66^.

To compare the differences in full-length versus partial genome hits across the six ESKAPE species, we analyzed the results generated from AMRFinderPlus. For each species, the proportions of full-length and partial genome hits were calculated for each sequencing method. We employed Pearson’s Chi-squared test with Yates’ continuity correction to assess the statistical significance of differences in these proportions. In addition, pairwise comparisons were made between each combination of sequencing methods (Illumina vs. ONT, Illumina vs. Hybrid, and ONT vs. Hybrid).

Finally, to assess the association between sequencing depth (log-transformed total number of bases sequenced) and the number of identified antimicrobial resistance (AMR) genes and point mutations, a linear regression was performed. This analysis aimed to evaluate the relationship between sequencing deepness and the number of detected genes across all samples. Subsequently, an analysis of variance (ANOVA) was conducted to compare the number of AMR genes and point mutations detected across different sequencing platforms.

Sequencing models were categorized into two groups: ONT (Oxford Nanopore Technologies), including MinION, GridION, and PromethION, and Illumina, including MiniSeq, MiSeq, NextSeq, NovaSeq, iSeq, and HiSeq. The number of detected AMR genes was the dependent variable, with sequencer model as the main factor and platform (Illumina and ONT) as a covariate. An interaction term between sequencer and platform was also included to evaluate whether the effect of sequencing models on gene detection varied based on the sequencing platform. All analyses were performed in R ^66^.

## Results

Overall, all the identified 1,385 sequences were used as no sequences were flagged as poor quality. From those sequences, 91 isolates belong to *E. faecium,* 201 to *S. aureus,* 537 to *K. pneumoniae,* 257 to *A. baumannii,* 159 to *P. aeruginosa,* and 140 to *E. cloacae*.

### Genome completeness

The assessment of genome completeness using BUSCO showed that isolates sequenced by Illumina can reach a higher level of completeness than those sequenced by ONT. The application of hybrid assembly reached similar completeness levels than those observed for Illumina (Table 1, Supplementary table 2).

**Table 1:**
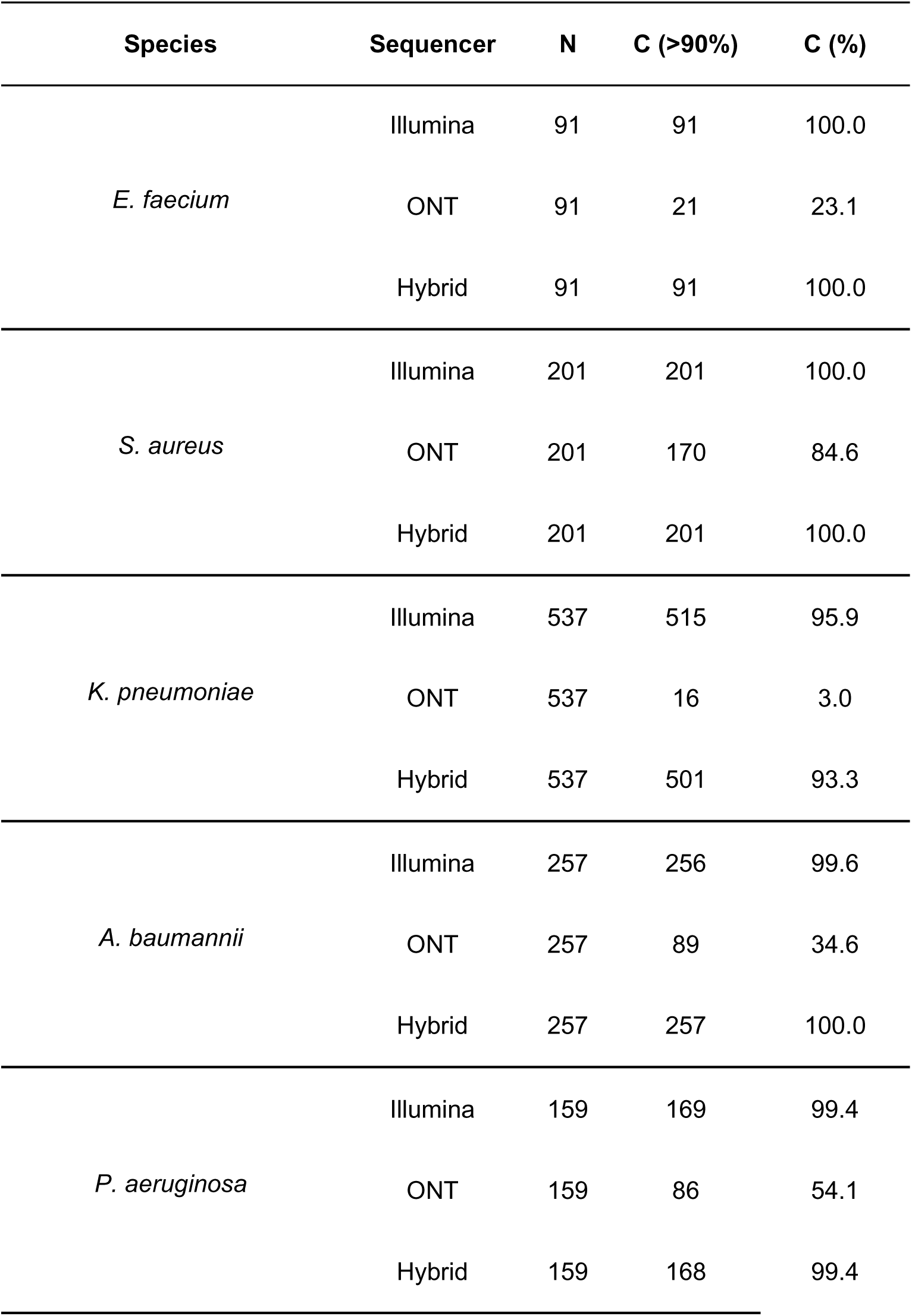
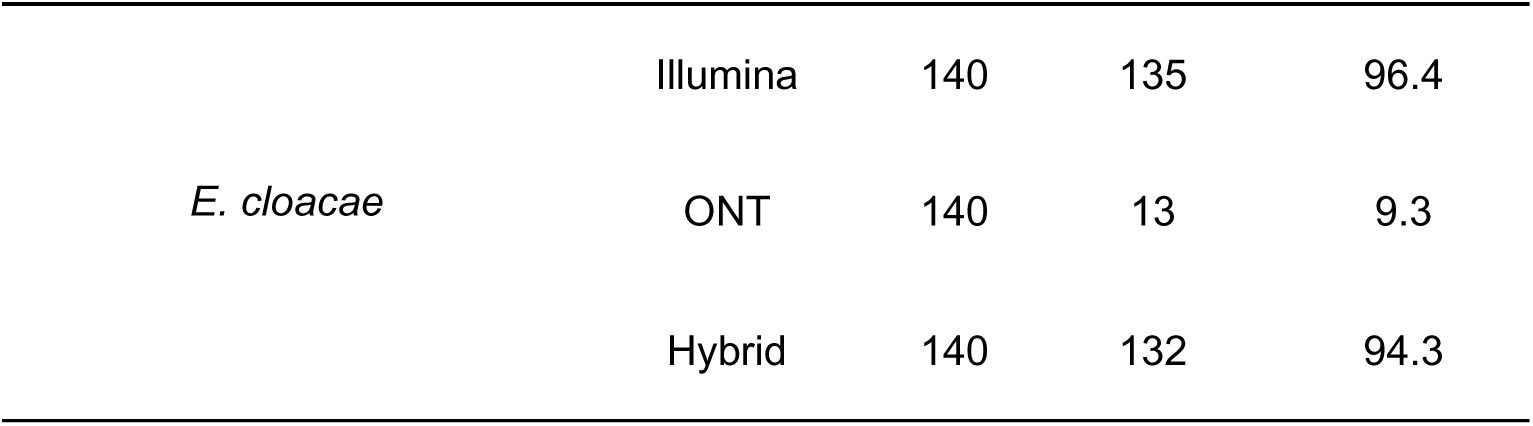
Genome completeness (Based on BUSCO results) determined on Illumina, Oxford Nanopore Technologies (ONT), and hybrid method on ESKAPE pathogens. C (>90%) corresponds to the number of sequences surpassing the 90% threshold of coverage as indicated by the completeness metric from BUSCO. C (%) indicates the percentage of the total number of sequences that surpassed that 90% threshold.

Overall, the results of the comparison of the differences in full-length versus partial genome hits show that Illumina sequencing provides reliable full-length hits with low percentage of partial hits. Even though ONT sequences presented high full-length genome coverage, they also showed a high proportion of partial hits. In contrast, the hybrid method offers a balanced approach, combining the strengths of both Illumina and ONT to achieve comprehensive and accurate identification of AMR genes (Supplementary Figure 1).

### AMR gene profiles across ESKAPE pathogens

Overall, we observed a wide diversity of resistance profiles among ESKAPE pathogens. All three methods (short-reads, long-reads, and hybrid assembly) detected a wide variety of resistance genes (detailed list available in Supplementary Table 2). The general trend across the six species consisted of Hybrid assembly detecting an overall higher number of AMR genes, followed by Illumina and ONT (Supplementary Table 2).

### Overall AMR gene and point mutation detection

We observed statistically significant differences (p<0.05) in AMR detection between the three methods (Figures 2-7 panel A; Supplementary Table 3). For all species, Hybrid assembly showed the highest detection power for AMR genes, outperforming both Illumina and ONT. The detection power of each technology is calculated by comparing the percentage of times each AMR gene or point mutation was detected by each technology (Supplementary Tables 2 and 3). β-lactam resistance emerged as the AMR class with the highest number of genes identified by all three sequencing methods following the same pattern where Hybrid assembly was the technology detecting a higher number of genes, followed by Illumina and ONT, except *for E. faecium* (0 genes detected) and *S. aureus* (7 genes), where Illumina detected more genes than both Hybrid assembly and ONT.

**Figure 2.**
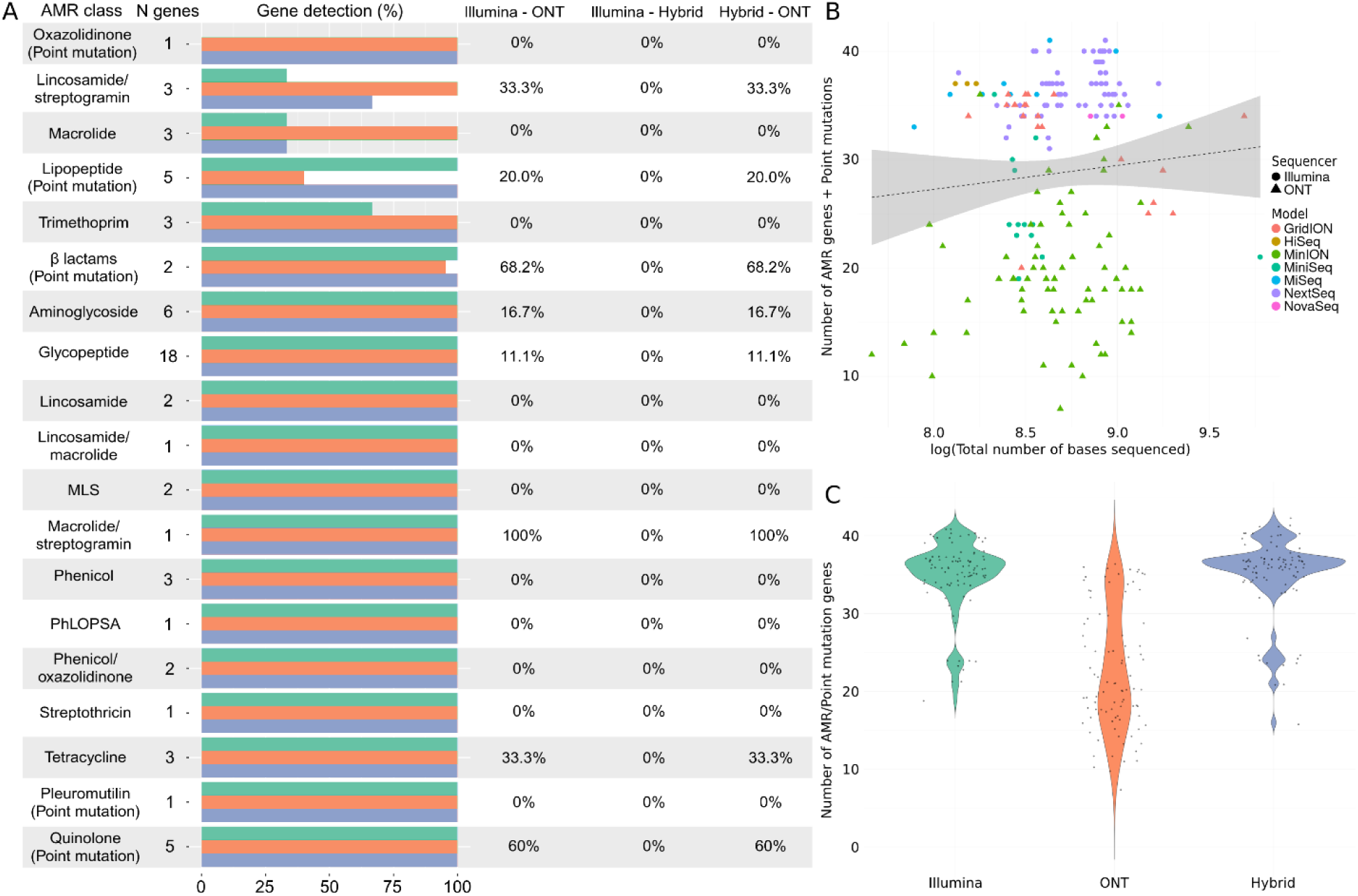
Comparative analysis of antimicrobial resistance detection in *E. faecium* using different sequencing technologies. A represents the differences in the identification of AMR genes and point mutations conferring resistance by Illumina (green), Oxford Nanopore Technologies (ONT) (orange), and hybrid assembly (blue). The horizontal bars represent the total number and its corresponding percentage of genes detected within each antimicrobial class, followed by the percentage of genes that were significantly differently detected by each pair of sequencers (0% = no differences in detection). B pictures the relationship between the total number of sequenced bases by each sequencer model, from Illumina (HiSeq, MiniSeq, MiSeq, NextSeq, and NovaSeq) and ONT (GridION, MinION and PromethION). C shows the overall differences in the number of AMR and point mutation genes detected by Illumina and ONT sequencing platforms, where the y-axis represents the total number of genes and point mutations identified.

When the overall detection of point mutations was compared, we observed that Illumina exhibited higher detection power compared to both Hybrid assembly and ONT except for *S. aureus*, where Hybrid assembly detected a higher number. Although Illumina was generally able to detect a higher number of point mutations, Hybrid correction performed comparably well in most species with minimal differences found in its detection rates (Supplementary Tables 2 and 3). When we combined the detected AMR genes with the AMR-associated point mutations and compared them with the total number of bases sequenced per isolate, we did not find a specific pattern across the species. *E. faecium* and *A. baumannii* showed positive correlations between sequencing depth and the number of AMR genes and point mutations. On the contrary, *S. aureus, K. pneumoniae, P. aeruginosa* and *E. cloacae* showed a negative correlation between these variables. To further explore these results, we also analyzed the potential differences between AMR gene and point mutation detection within each sequencing technology by analyzing each platform as a covariate in ANCOVA analyses. These results evidence the influence of sequencing depth in all species except for *S. aureus,* where these results were these differences were not statistically significant. Similarly, when we compared the overall detection of AMR genes and point mutations by each sequencing method, all species except for *S. aureus* showed significant differences, where Hybrid correction identified the highest number, closely followed by Illumina and finally by ONT, which generally detected a significantly smaller number.

### ESKAPE specific AMR profiles

We found significant differences in AMR gene detection between the three methods, with Hybrid correction detecting a higher number of genes, followed by Illumina and ONT, which identified a significantly fewer number (Figure 2 panel A, Supplementary Table 3). Some examples of these differences were particularly clear in the case of aminoglycosides, glycopeptides, lincosamide/streptogramins and tetracyclines (Figure 2, Supplementary Tables 2 and 3).

In the case of point mutations, the isolates sequenced by Illumina exhibited the highest number, followed by hybrid assembly and finally by ONT (Supplementary Table 3). Overall, the highest number of point mutations detected were associated with β-lactam genes, followed by lipopeptide and quinolone-associated genes (Figure 2, Supplementary Table 3). We found no significant association between sequencing depth of all *E. faecium* isolates used in this study and the overall combination of AMR genes plus point mutations (*p* > 0.05). However, we did not find significant differences between sequencer models (i.e. MiSeq, NovaSeq, MinION, GridION, etc) when the sequencing technology was analyzed as a covariate (Illumina or ONT) following a positive association between number of genes and point mutations and sequencing depth (Figure 2, panel B, Supplementary Table 4, ANCOVA results). Similarly, we did not find statistically significant differences in the overall number of AMR genes plus point mutations combined identified by each method (Hybrid, Illumina, ONT) (Figure 2, panel C, Supplementary Table 4 ANOVA results), with Hybrid correction detecting the highest number.

The results for *S. aureus* also showed that overall, hybrid assembly found the highest number of AMR genes, followed by Illumina and ONT (Supplementary Tables 2 and 3). These differences were significant for a number of antimicrobial classes such as β-lactams or glycopeptides (particularly the *Van* gene family). In the case of point mutations, Hybrid assembly was able to detect a higher number than Illumina and ONT. These mutations were particularly present in genes associated with fosfomycin, quinolone and sulfonamide resistance (Figure 3 panel A, Supplementary Tables 2 and 3). When comparing the relationship between sequencing depth of all *S. aureus* isolates used in this study and the total number of AMR genes and point mutations, we found no significant differences (Figure 3 panel B, Supplementary Table 4). Likewise, there were no differences on the overall number of AMR genes and point mutations identified, showing a consistent pattern across the three sequencing methods (Figure 3, panel C, Supplementary Table 4).

**Figure 3.**
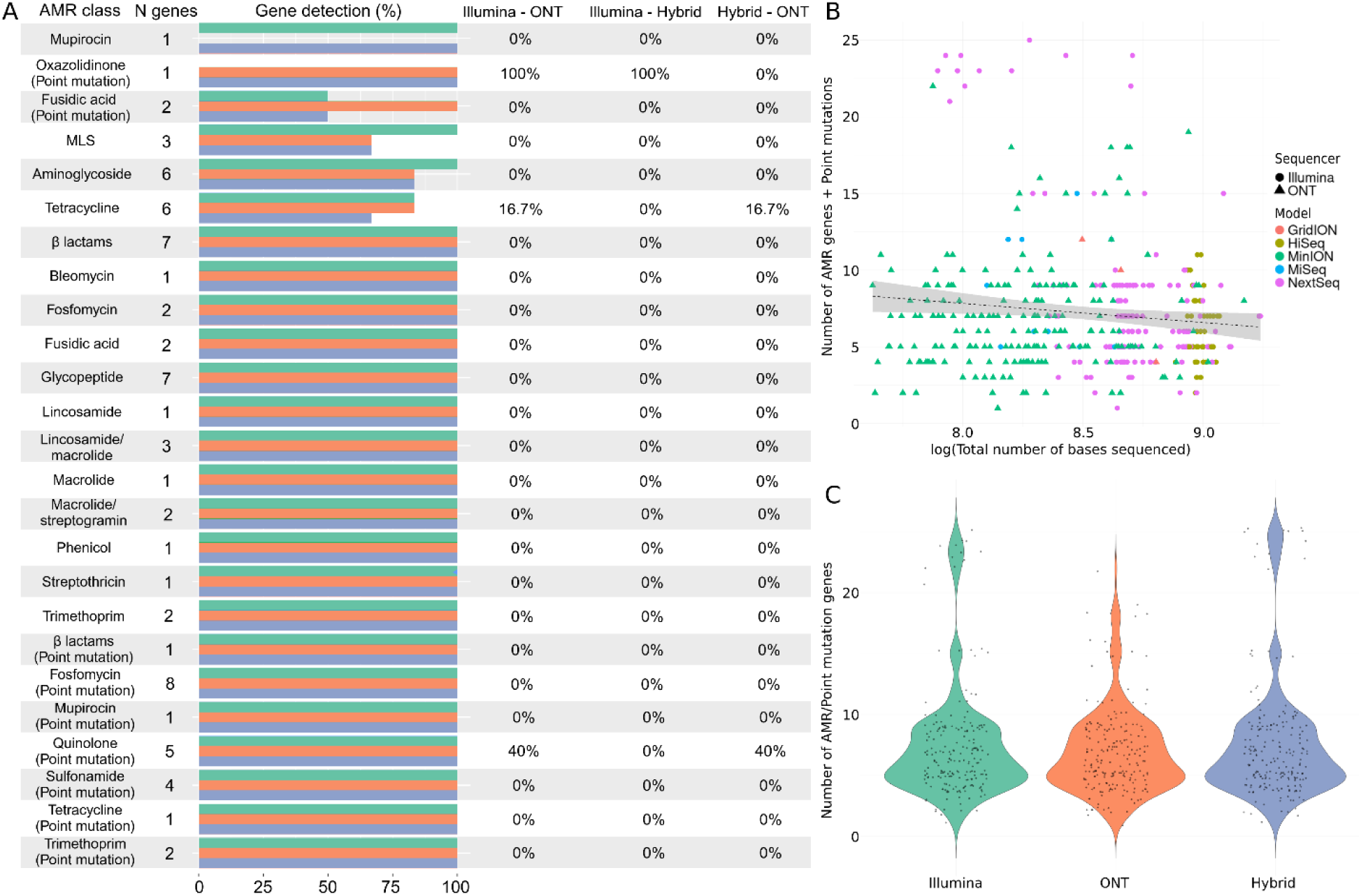
Comparative analysis of antimicrobial resistance detection in *S. aureus* using different sequencing technologies. A represents the differences in the identification of AMR genes and point mutations conferring resistance by Illumina (green), Oxford Nanopore Technologies (ONT) (orange), and hybrid assembly (blue). The horizontal bars represent the total number and its corresponding percentage of genes detected within each antimicrobial class, followed by the percentage of genes that were significantly differently detected by each pair of sequencers (0% = no differences in detection). B pictures the relationship between the total number of sequenced bases by each sequencer model, from Illumina (HiSeq, MiniSeq, MiSeq, NextSeq, and NovaSeq) and ONT (GridION, MinION and PromethION). C shows the overall differences in the number of AMR and point mutation genes detected by Illumina and ONT sequencing platforms, where the y-axis represents the total number of genes and point mutations identified.

We observed a wide variety of genes that showed significant differences in the detection of AMR genes depending on the sequencing method used for *K. pneumoniae* sequences. It is important to note that *K. pneumoniae* was the species with the highest number of analyzed isolates of the six, which likely contributed to these results.

*K. pneumoniae* followed the same pattern as *E. faecium*, where Hybrid assembly detected the highest number of AMR genes followed by Illumina, with ONT identifying significantly fewer genes (Figure 4 panel A, Supplementary Tables 2 and 3). Genes associated to carbapenem resistance (β-lactam class) presented the largest discrepancies between the sequencing methods, where Hybrid assembly and Illumina showed similar levels of detection, both outperforming ONT. This same pattern was also observed for quinolone and aminoglycoside resistance, where ONT sequences identified significantly fewer genes than the other two methods. Hybrid assembly also showed a slightly higher detection of point mutations than Illumina, followed by ONT which detected significantly fewer (Supplementary Table 3). These mutations were more often detected for β-lactam, quinolone and tetracycline genes (Figure 4, Supplementary Tables 2 and 3).

**Figure 4.**
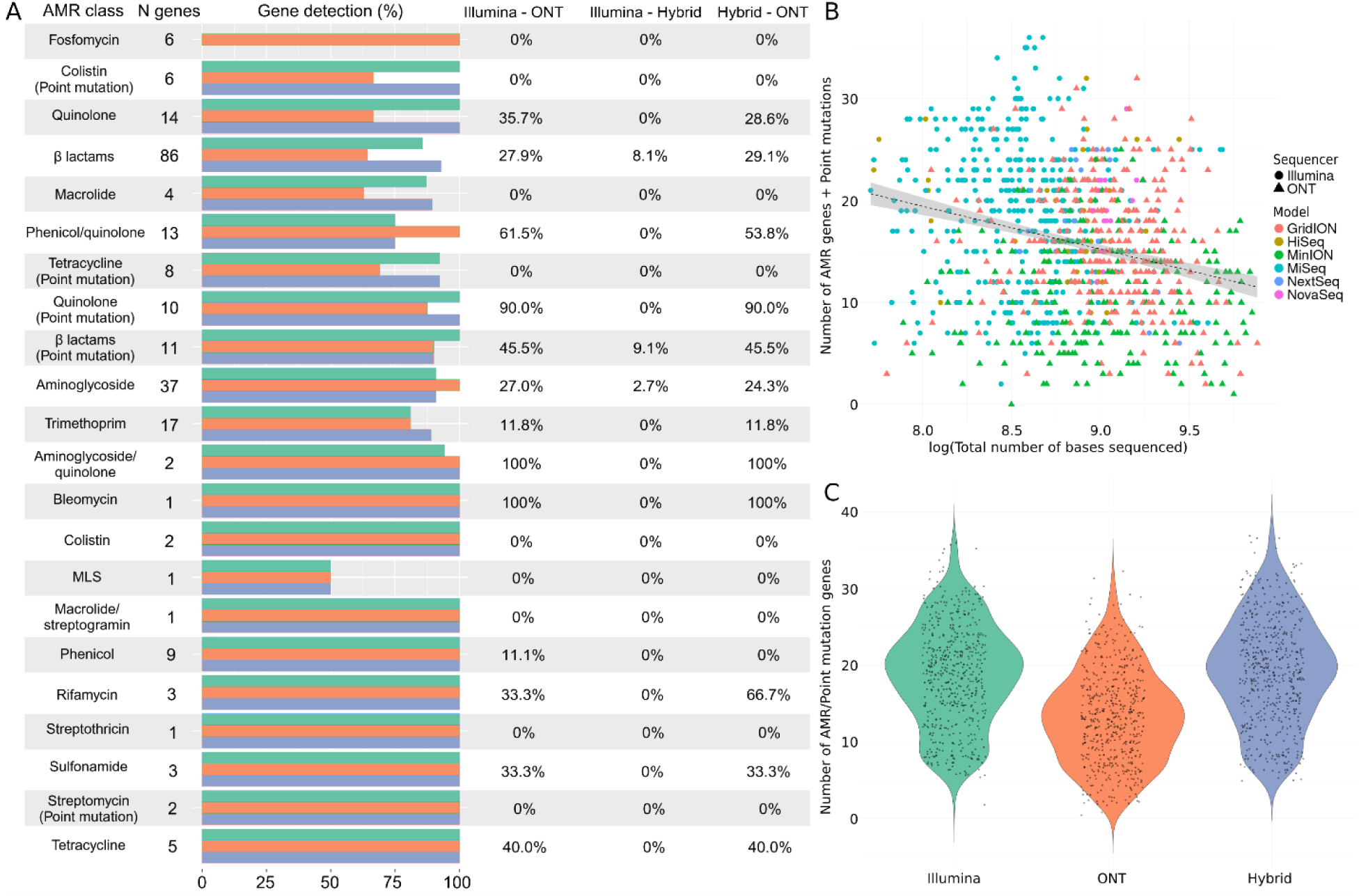
Comparative analysis of antimicrobial resistance detection in *K. pneumoniae* using different sequencing technologies. A represents the differences in the identification of AMR genes and point mutations conferring resistance by Illumina (green), Oxford Nanopore Technologies (ONT) (orange), and hybrid assembly (blue). The horizontal bars represent the total number and its corresponding percentage of genes detected within each antimicrobial class, followed by the percentage of genes that were significantly differently detected by each pair of sequencers (0% = no differences in detection). B pictures the relationship between the total number of sequenced bases by each sequencer model, from Illumina (HiSeq, MiniSeq, MiSeq, NextSeq, and NovaSeq) and ONT (GridION, MinION and PromethION). C shows the overall differences in the number of AMR and point mutation genes detected by Illumina and ONT sequencing platforms, where the y-axis represents the total number of genes and point mutations identified.

*K. pneumoniae* presented a strong significant negative relationship between the total number of bases sequenced of all isolates and the number of antimicrobial resistance genes and point mutations identified. These differences were observed both between sequencing technologies (Illumina vs ONT) and also when these technologies were assessed as a covariate (Figure 4 panel B, Supplementary Table 4). Similarly, we found significant differences between the number of AMR genes and point mutations identified by each of the methods, with Hybrid assembly detecting a higher number than Illumina, followed by ONT (Figure 4 panel C, Supplementary Table 4).

In the case of *A. baumannii*, Hybrid assembly detected a slightly higher number of AMR genes, followed by Illumina and ONT. The difference between ONT with the other two approaches was particularly significant for antimicrobial classes such as β-lactams and aminoglycosides, which were the two most common in this species (Figure 5 panel A, Supplementary Tables 2 and 3). In the case of point mutations, both Hybrid assembly and Illumina identified an equal number of mutations, with ONT detecting significantly fewer (Supplementary Tables 2 and 3). A positive correlation between sequencing depth and the number of identified AMR genes and point mutations was observed in *A. baumannii* sequences both between platforms (Illumina vs ONT) and when each sequencing technology was analyzed as a covariate although the correlation was weak (Figure 5 panel B, Supplementary Table 4). Additionally, slight but significant differences were detected in the number of AMR genes identified by each method, with Hybrid assembly detecting a higher number than Illumina or ONT (Figure 5, panel C, Supplementary Table 4).

**Figure 5.**
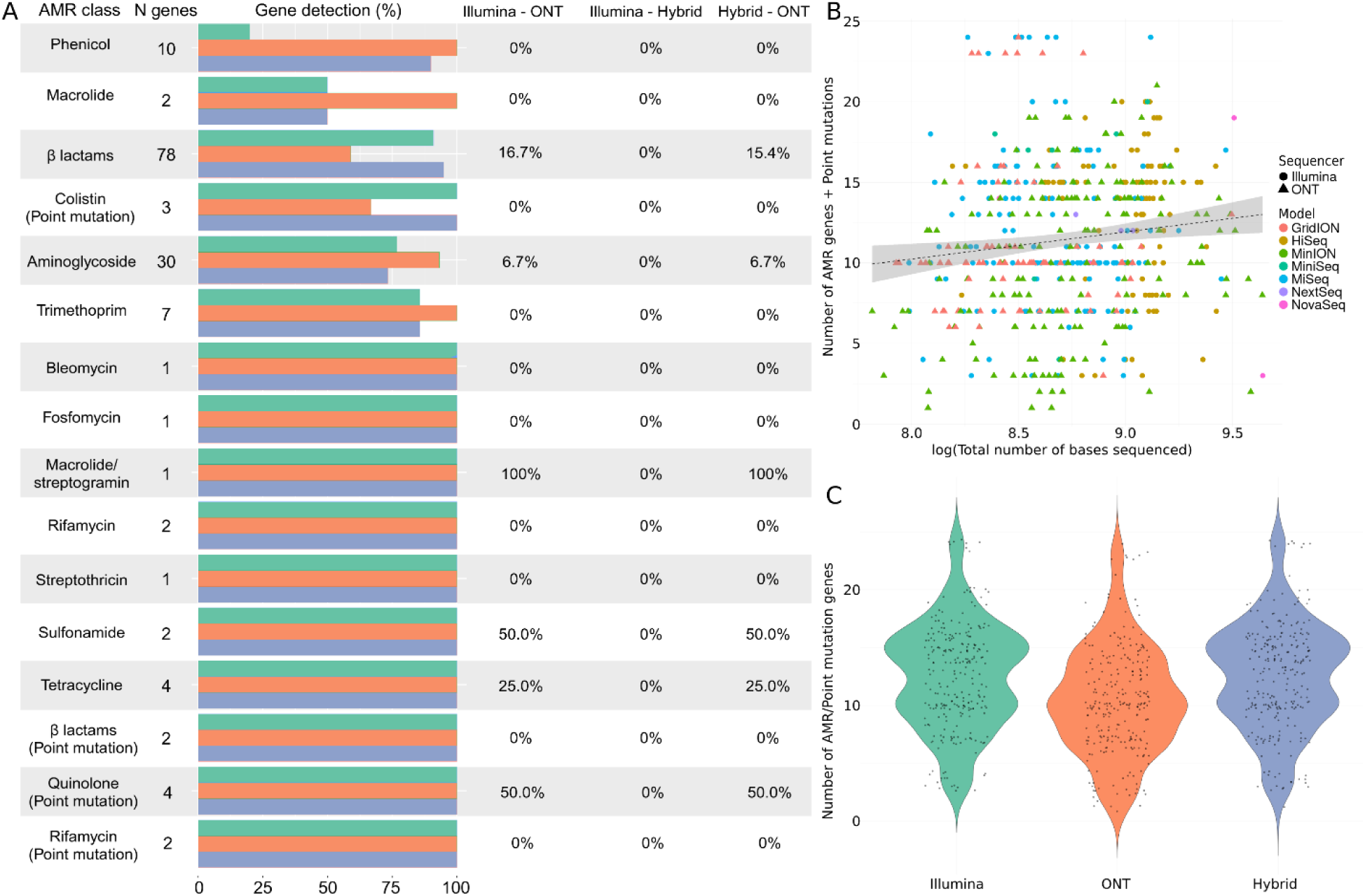
Comparative analysis of antimicrobial resistance detection in *A. baumannii* using different sequencing technologies. A represents the differences in the identification of AMR genes and point mutations conferring resistance by Illumina (green), Oxford Nanopore Technologies (ONT) (orange), and hybrid assembly (blue). The horizontal bars represent the total number and its corresponding percentage of genes detected within each antimicrobial class, followed by the percentage of genes that were significantly differently detected by each pair of sequencers (0% = no differences in detection). B pictures the relationship between the total number of sequenced bases by each sequencer model, from Illumina (HiSeq, MiniSeq, MiSeq, NextSeq, and NovaSeq) and ONT (GridION, MinION and PromethION). C shows the overall differences in the number of AMR and point mutation genes detected by Illumina and ONT sequencing platforms, where the y-axis represents the total number of genes and point mutations identified.

The detection of AMR genes for *P. aeruginosa* was higher for Hybrid assembly, closely followed by Illumina and with ONT detecting fewer genes overall. Aminoglycoside and β-lactam resistance genes were the most prevalent, with the *aac* and *bla* gene families as the most detected (Figure 6 panel A, Supplementary Tables 2 and 3). Similarly, point mutations associated with β-lactams were the most commonly found in this bacterium, followed by mutations associated with quinolones. As in other species, Illumina detected the highest number of point mutations, closely followed by hybrid assembly and ONT. There were no statistically significant associations between the overall sequencing depth of *P. aeruginosa* isolates and the number of antimicrobial resistance genes identified in *P. aeruginosa*. However, there was a significant influence of sequencing deepness when sequencing platforms were measured as a covariate, showing a positive correlation (Figure 6, panel B, Supplementary Table 4). Likewise, our results showed significant differences in the overall detection of AMR genes and point mutations between the three methods, with Hybrid assembly identifying the highest number (Figure 6 panel C, Supplementary Table 4).

**Figure 6.**
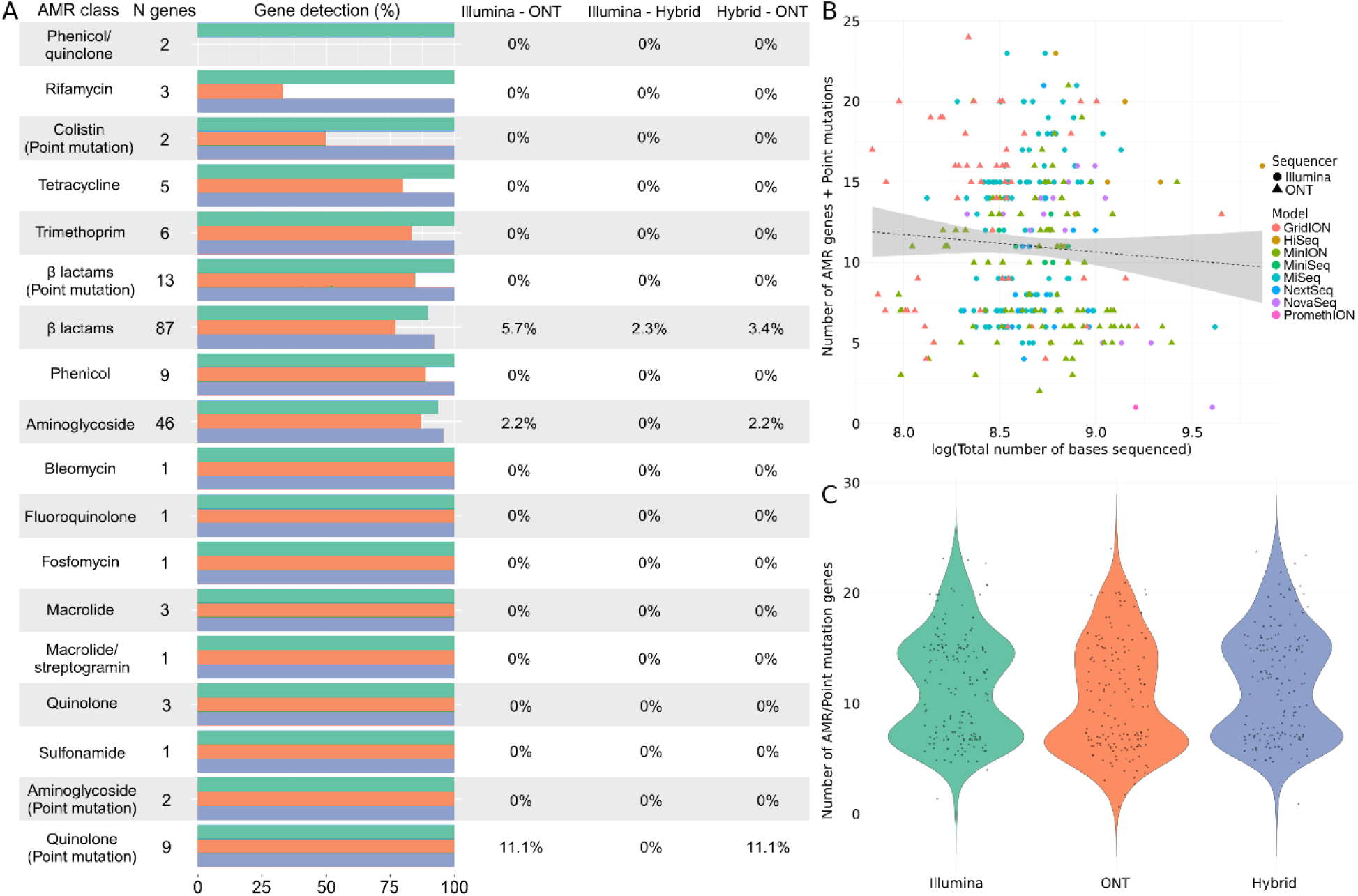
Comparative analysis of antimicrobial resistance detection in *P. aeruginosa* using different sequencing technologies. A represents the differences in the identification of AMR genes and point mutations conferring resistance by Illumina (green), Oxford Nanopore Technologies (ONT) (orange), and hybrid assembly (blue). The horizontal bars represent the total number and its corresponding percentage of genes detected within each antimicrobial class, followed by the percentage of genes that were significantly differently detected by each pair of sequencers (0% = no differences in detection). B pictures the relationship between the total number of sequenced bases by each sequencer model, from Illumina (HiSeq, MiniSeq, MiSeq, NextSeq, and NovaSeq) and ONT (GridION, MinION and PromethION). C shows the overall differences in the number of AMR and point mutation genes detected by Illumina and ONT sequencing platforms, where the y-axis represents the total number of genes and point mutations identified.

Finally, *E. cloacae* presented significant differences in AMR gene detection between the methods, following the same pattern described previously, where Hybrid correction detected a higher number of genes, followed by Illumina and finally ONT. Genes associated with aminoglycosides and β-lactams were the most prevalent, with the *aac* and *bla* gene families as the main groups of genes respectively (Figure 7 panel A, Supplementary Tables 2 and 3). *E. cloacae* only showed point mutations associated with quinolone resistance, particularly in the *gyrA* genes (Supplementary Tables 2 and 3), with Illumina detecting a higher number, followed by Hybrid assembly and ONT. *E. cloacae* presented a significant negative relationship between the total number of bases sequenced and the number of identified AMR genes and point mutations both when all sequences were analyzed together and when each platform was analyzed as a covariate (Figure 7, panel B, Supplementary Table 4). This bacterium also showed significant differences between the number of AMR genes and point mutations identified by the three methods, with hybrid assembly detecting the highest number, followed by Illumina and ONT (Figure 7, panel C, Supplementary Table 4).

**Figure 7.**
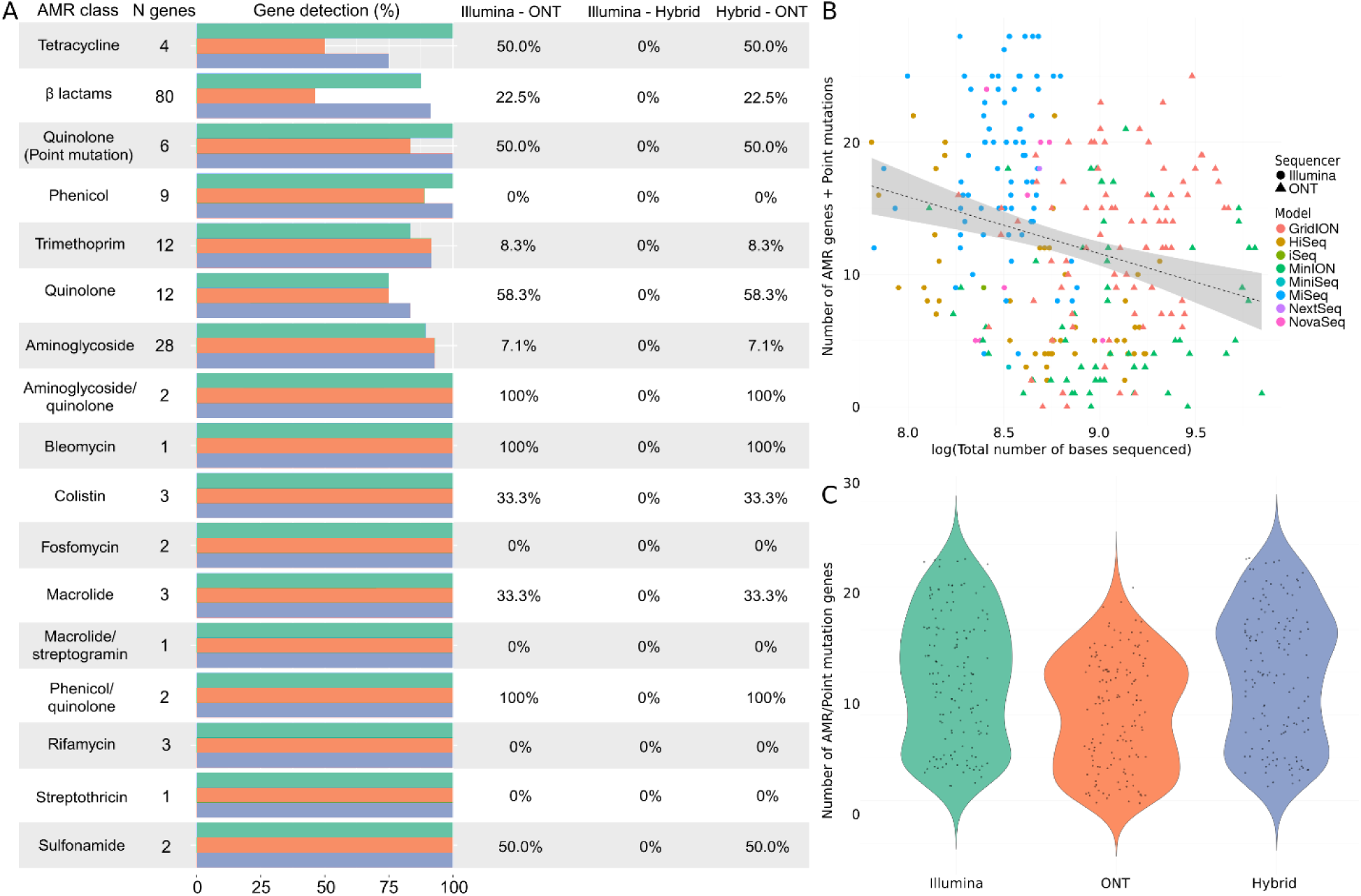
Comparative analysis of antimicrobial resistance detection in *E. cloacae* using different sequencing technologies. A represents the differences in the identification of AMR genes and point mutations conferring resistance by Illumina (green), Oxford Nanopore Technologies (ONT) (orange), and hybrid assembly (blue). The horizontal bars represent the total number and its corresponding percentage of genes detected within each antimicrobial class, followed by the percentage of genes that were significantly differently detected by each pair of sequencers (0% = no differences in detection). B pictures the relationship between the total number of sequenced bases by each sequencer model, from Illumina (HiSeq, MiniSeq, MiSeq, NextSeq, and NovaSeq) and ONT (GridION, MinION and PromethION). C shows the overall differences in the number of AMR and point mutation genes detected by Illumina and ONT sequencing platforms, where the y-axis represents the total number of genes and point mutations identified.

### Pangenome characterization

Generally, a consistent trend emerged in identifying genes across all six species. Among the three sequencing methods employed, Illumina-sequenced isolates consistently exhibited the lowest number of detected genes, while ONT-sequenced counterparts demonstrated the highest count. The hybrid assembly method typically yielded gene counts falling between those obtained from Illumina and ONT. This pattern can be observed in panel A of Supplementary Figures 2-7, which visualizes the comparison of the number of genes in the core genome detected by each technology (Illumina, ONT, and hybrid correction).

However, species-specific proportions of cloud, core, soft-core, and shell genes did not exhibit discernible patterns associated with the sequencing type but with the bacterial species (Table 2). The proportion of genes per pangenome structure (core, soft-core, shell, and cloud) detected by each method is available in panel B of Supplementary Figures 2-7. A recurrent observation across the species was that Illumina sequencing consistently revealed fewer unique accessory genes than the other methods. This can be observed by looking at the pangenome rarefactions curves (Supplementary Figures 2-7 panel C). Pangenome rarefaction curves are graphical visualizations of the growth of the pangenome size as a function of the number of genomes (or isolates) that are included in the analysis. Each point indicates the number of genes present in a specific genome. Therefore, genes that are not identical between two samples are identified as two individual genes, even if they are the same gene variants. As the number of analyzed genomes increases, so does the value found in the x-axis.

**Table 2:**
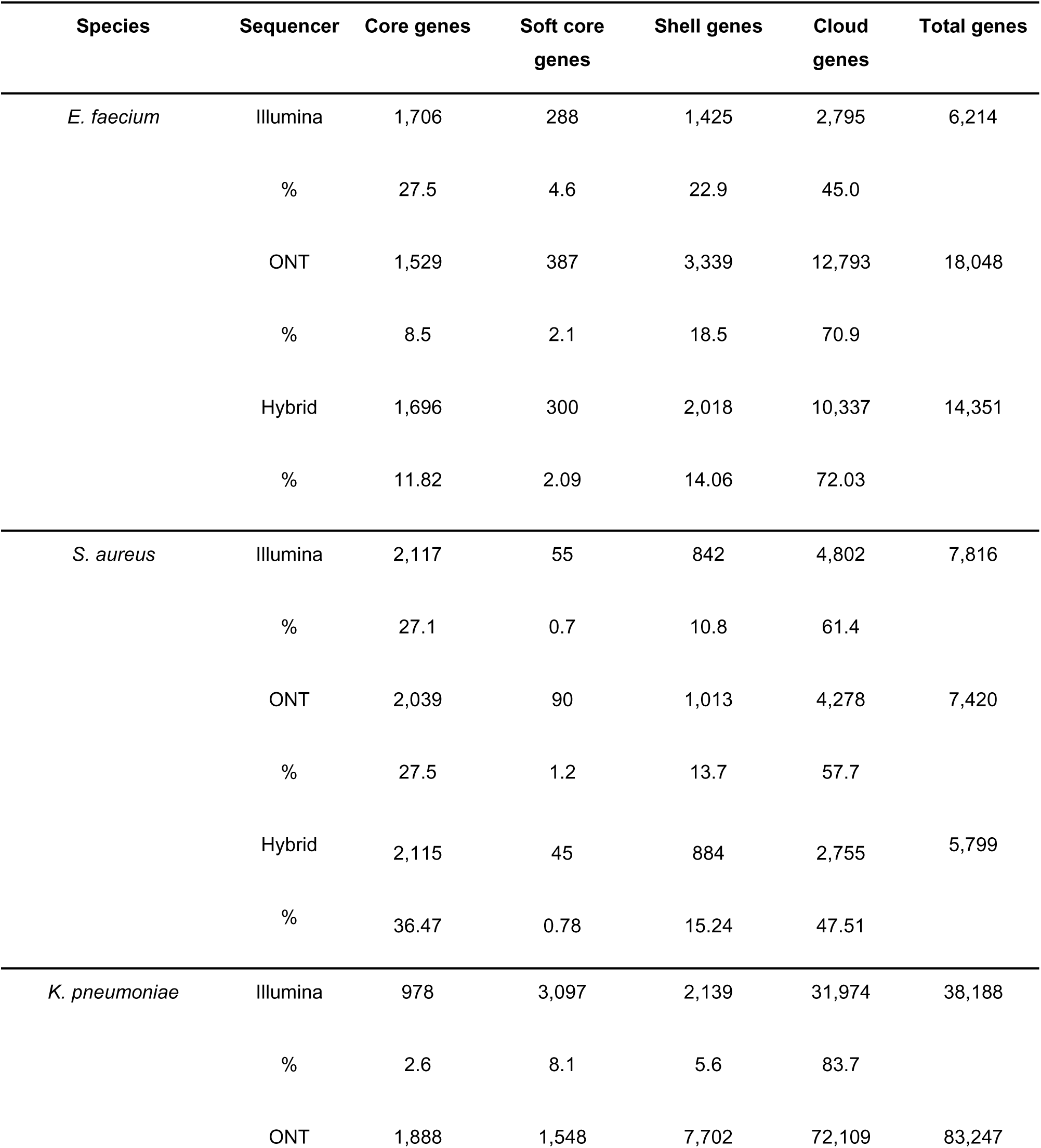
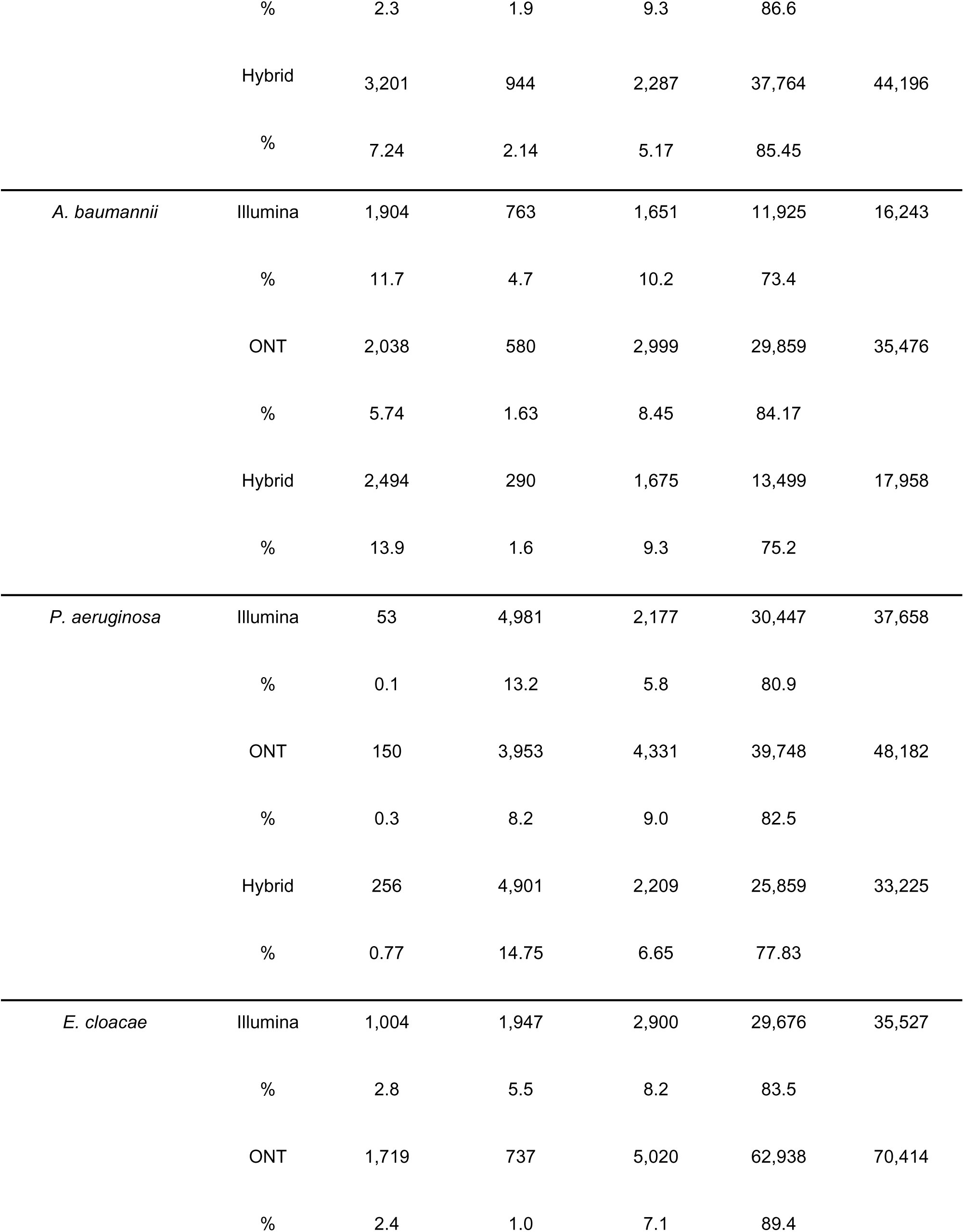
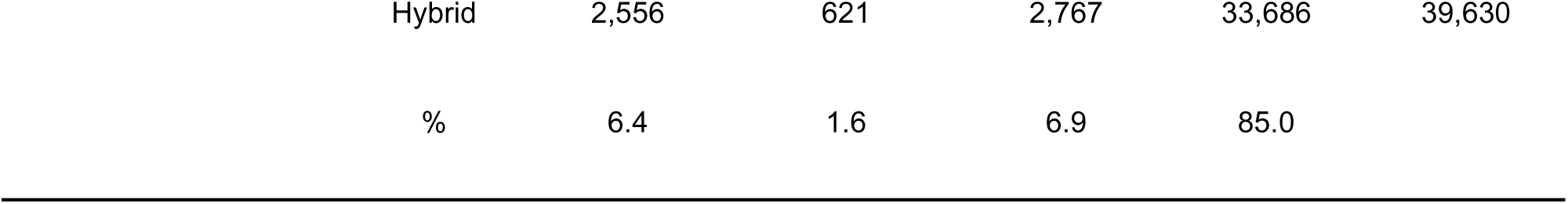
Summary of the pangenome gene classification in ESKAPE pathogens.

The shape of these curves represents the open or closed nature of the pangenome, where a continuously rising curve corresponds to an open pangenome meaning that as new samples are included a significant number of genes are added to the pangenome. On the other hand, if the rarefaction curve reaches a plateau, it indicates a closed pangenome since adding new isolates does not increase the pangenome size. In our case, Illumina sequences tend to lead to a narrower shape of the rarefaction curves, showing that this technology tends to find more genomic similarities and, therefore, fewer unique gene variants between the isolates (blue curve in panel C in Supplementary Figures 2-7). This suggests that Illumina sequences tend to identify a less diverse group of isolates within each species, except for *P. aeruginosa*, where this particular pattern was not observed.

For *E. faecium*, ONT identified the highest number of core genes, followed by hybrid correction and Illumina (panel A, Supplementary Figure 2), which exhibited a reduced proportion of accessory genome compared to ONT or hybrid assembly. The latter displayed a pattern more similar to ONT results (panel B, Supplementary Figure 2). These distinctions are observable in the pangenome rarefaction curves, where ONT and hybrid assembly reveal a higher share of distinctive accessory genes (panel C Supplementary Figure 2).

Conversely, *S. aureus* isolates sequenced using Illumina showed a closer number of core genes to those detected by ONT, with most genes detected by hybrid correction being shared between the methods (panel A, Supplementary Figure 3). Following this pattern, Illumina presented a more similar percentage of cloud genes than ONT and hybrid correction, with hybrid correction showing the lowest proportion of cloud genes of the three (Supplementary Figure 3). Finally, this bacterium’s rarefaction curves were more similar, with ONT still presenting a slightly wider shape (panel C, Supplementary Figure 3).

For *K. pneumoniae*, ONT detected a distinctively higher number of core genes compared to the other two methods, which were more similar when compared (panel A, Supplementary Figure 4). There were no significant variations in the proportion of cloud genes identified by each method. Nevertheless, Illumina exhibited the greatest ratio of soft core genes, ONT displayed the highest percentage of shell genes, and with hybrid correction, there was a comparable distribution between core and shell genes, while the number of soft core genes was the lowest among all methods (panel B, Supplementary Figure 4). This was observed in the rarefaction curves, where hybrid correction displayed the widest shape of the three (panel C, Supplementary Figure 4).

The findings for *A. baumannii* resemble those of *E. faecium* and *K. pneumoniae*. ONT revealed the highest number of core genes (panel A, Supplementary Figure 5), and the highest percentage of cloud genes, while the percentage of shell genes remained consistent across all three options. Ultimately, Illumina displayed the highest proportion of soft-core genes (panel A, Supplementary Figure 5). The shape of the rarefaction curves was more similar between Illumina and hybrid correction, with ONT presenting a slightly wider distribution (panel C, Supplementary Figure 5).

*P. aeruginosa* stood out as the sole species with an almost negligible core genome (0.1-0.7%), showing a clear scattered pangenome curve when compared to the other species. For this bacterium, the number of core genes was similar between the three methods, with ONT still detecting a higher number of genes (panel A, Supplementary Figure 6). Hybrid correction yielded fewer cloud genes, whereas Illumina and ONT exhibited comparable percentages. Regarding shell genes, ONT sequences showed the highest proportion, followed by the hybrid correction, and Illumina. Notably, the proportion of soft-core genes was similar between Illumina and hybrid correction, both surpassing the levels detected in ONT (panel B, Supplementary Figure 6). The rarefaction curves for this species were generally spread in shape, with Illumina being the wider of the three (panel C, Supplementary Figure 6).

*E. cloacae* followed similar patterns to *K. pneumoniae,* with ONT detecting the highest number of core genes compared to either Illumina or hybrid correction (panel A, Supplementary Figure 7). A comparable proportion of shell genes was identified by all three methods. Illumina sequences exhibited the highest percentage of soft-core genes, whereas hybrid correction showed the highest percentage of core genes. Ultimately, ONT and hybrid approach identified the highest proportion of cloud genes (panel B, Supplementary Figure 7). Finally, the shape of the rarefaction curves for this species was similar between the three methods, with hybrid correction presenting a slightly wider distribution (panel C, Supplementary Figure 7).

The results of the ANOVA tests comparing the number of annotated genes prior to pangenome reconstruction showed statistically significant differences between the three approaches (Illumina, ONT, and hybrid assembly) across all six bacterial species. Illumina sequences consistently presented fewer annotated genes compared to both ONT and hybrid assembly, but these two did not show statistical differences between each other in any of the bacteria (Supplementary Table 5, Supplementary Figure 8).

## Discussion

This comprehensive study was designed to compare the outcomes of similar bioinformatic analyses conducted on isolates from ESKAPE pathogens (*Enterococcus faecium, Staphylococcus aureus, Klebsiella pneumoniae, Acinetobacter baumannii, Pseudomonas aeruginosa,* and *Enterobacter cloacae*) that were sequenced with both short-read (Illumina) and long-read (Oxford Nanopore Technologies, ONT) platforms, as well as the emerging hybrid assembly approach. While Illumina and ONT sequencing effectively identified key AMR genes, our findings revealed significant variability across the different methods.

The differences in performance between ONT and Illumina can be partially attributed to the fact that most bioinformatic tools have historically been developed to process short-read sequences due to the longer presence and accessibility of Illumina technology ^67, 68^. Consequently, these tools have had more time to refine their pipelines and algorithms for short-read data analysis, which may not be optimized for the distinct characteristics of ONT long-read sequences. This disadvantage was observed in our genome completeness (BUSCO) results, which showed better performance on Illumina sequences compared to ONT. Despite discrepancies in our BUSCO results, we included these findings to provide complementary information supporting the observed differences between Illumina and ONT sequencers. BUSCO scoring of predicted genes is based on Hidden Markov Model (HMM) profiles that must meet a score threshold ^69^. BUSCO’s gene predictor can encounter spurious stop codons caused by the high error rate of the long-read technology. As a result, a truncated protein is passed to HMM to be scored, which will lower BUSCO results. With Illumina, the error rate is lower, so these early stop codons are not as common (Mathieu Seppey personal communication, BUSCO developer). We observed a substantial improvement in assessing genome completeness when applying the hybrid assembly method, which also led to an increase in the number of AMR genes identified, nearing what we found in the isolates sequenced with Illumina. These results demonstrate how discrepancies in tool development timelines underscore the need for further advancements in bioinformatics tailored to long-read data analysis to fully harness the potential of ONT technology in genomics and molecular research, highlighting the need for improved tools for long-reads.

Our results align with the previously described AMR repertoire for ESKAPE pathogens, with a high prevalence of genes conferring resistance to β-lactams throughout most species ^70^. β-lactam genes are known to present a wide variety of closely related variants ^71, 72^, where short-read sequencing tends to have higher accuracy at individual-base level than long-read sequencing, due to its better performance for single nucleotide polymorphism (SNP) analysis ^13, 73^. The existence of these variants is likely the main reason behind the difference in the number of observed β-lactam genes between Illumina and ONT sequences, especially in species known to present a higher number of β-lactam gene variants such as *A. baumannii, E. cloacae*, and *P. aeruginosa*, the first two known for their complex resistomes ^74–76^. Genes conferring resistance against aminoglycosides, macrolides, and tetracyclines were consistently identified by both sequencers across all species, and therefore also by hybrid correction. This similarity indicates that current tools and databases for detecting AMR genes perform well across platforms. Resistance against other antimicrobial classes were more variable, independently of the sequencing method.

Similarly, our results also align with previous studies describing that Illumina sequencers tend to outperform ONT sequencers at point mutation detection in other organisms ^23, 77^. These differences primarily arise from the inherent error profiles associated with read length of each sequencing platform, positioning emerging approaches such as hybrid assembly as a great option to improve ONT accuracy while more robust downstream analysis pipelines are established ^31^.

We did not observe a conserved pattern showing the directionality of the relationship between overall sequencing depth and AMR gene and point mutation detection. However, the influence of sequencing deepness was evident in all species except for *S. aureus*, which presented comparable results across the three methods. These similarities could be due to the clinical importance of *S. aureus* and the detailed characterization of its genome, but also due to a potential higher similarity between the isolates used for the analyses.

This difference between the species was also observed in the characterization of the pangenome size and its composition, where *S. aureus* presented minimal variation between the three methods compared to the rest of ESKAPE pathogens. These differences most likely arise from variations in gene annotation, where isolates sequenced by Illumina consistently resulted in significantly fewer annotated genes compared to either ONT or hybrid assembly. These results indicate that the choice of sequencing technology can have a significant influence in gene annotation in bacterial genome assemblies, which may have potential implications for downstream analyses as pangenome reconstruction.

Thus, our results highlight that short and long-read sequencing technologies at their current state can present enough variations in the obtained outcomes that the decision of using either long or short sequences, or their combination through hybrid assembly should be driven by the specific goals, bacterial species, genes of interest and scope of the study. For instance, our results support previous literature that suggests Illumina as a great option for common AMR analysis and pangenome reconstruction, due to its per-base accuracy, which is crucial for confidently identifying resistance ^78^, while ONT can be a better option for the study of newly discovered bacterial species with potentially rearranged genomes since it is able to generate reads spanning tens of kilobases, offering a clearer view of large insertions or deletions, which in the case of pangenome reconstruction, will provide a comprehensive coverage of complex regions ^79, 80^. Our observations on the application of hybrid assembly, also prove its potential as a promising strategy to optimize the benefits of both technologies for a more comprehensive understanding of bacterial pathogens’ genetic makeup ^38, 81^.

Finally, it is noteworthy that our analyses were based on isolates for which both short and long-read sequences were already available, which may be an indicator of specific objectives on their original studies that could potentially bring bias of the type of isolates collected. For that reason, these isolates may not be accurate representations of the actual variation and diversity of the resistome in ESKAPE pathogens. One likely example of this is the unusual low resistance found for *S. aureus* isolates compared to previously described information. Therefore, the results of the current study must be interpreted aiming to identify and understand the differences that can be encountered in the obtained results depending on the use of short reads (Illumina), long reads (ONT), or hybrid assembly for the study of AMR, AMR-associated point mutations or pangenome composition and structure, and serve as a guide for a data-driven decision for future research.

## Conclusions

Our study compared short-read (Illumina) and long-read (ONT) sequencing technologies and the application of hybrid assembly, for analyzing AMR-related gene detection in ESKAPE pathogens at pangenome level. Both technologies identified key genetic elements, Hybrid assembly often outperformed both individual methods for AMR gene detection, closely followed by Illumina and finally by ONT, which detected significantly fewer genes. However, in the case of point mutation detection, Illumina was able to detect a higher number than the other two methods. Genome completeness was higher for short-read sequences, likely due to the existence of optimized bioinformatic tools.

While no consistent pattern of correlation between AMR detection and sequencing depth emerged across species, this variable significantly influenced results within individual platforms. Pangenome analysis showed that ONT can identify a higher number of core and accessory genes, while Illumina found fewer unique variants between isolates likely due to the consistently lower number of annotated genes detected by it. For these reasons, this study highlights the need to take both pathogen and specific goals of the study into consideration when selecting a sequencing approach. This study provides valuable insights into the strengths, limitations, and synergies of different sequencing technologies, guiding researchers in making informed choices for comprehensive genetic analyses of bacterial pathogens.

## Authors’ contributions

AFDD: conception and design of work, acquisition, analysis, and interpretation of data, writing, and revision of manuscript. MJ: conception and design of work, acquisition, analysis, and interpretation of data, writing, and revision of manuscript. CL: conception and design of work, interpretation of data, writing, and revision of manuscript, acquisition of funding, and supervision of project. All authors have approved the submitted version of the manuscript and have agreed to be accountable for their contribution to the article.

