## Supplementary Figures for "Influence of Sequencing Technology on Pangenome-level Analysis and Detection of Antimicrobial Resistance Genes in ESKAPE Pathogens"

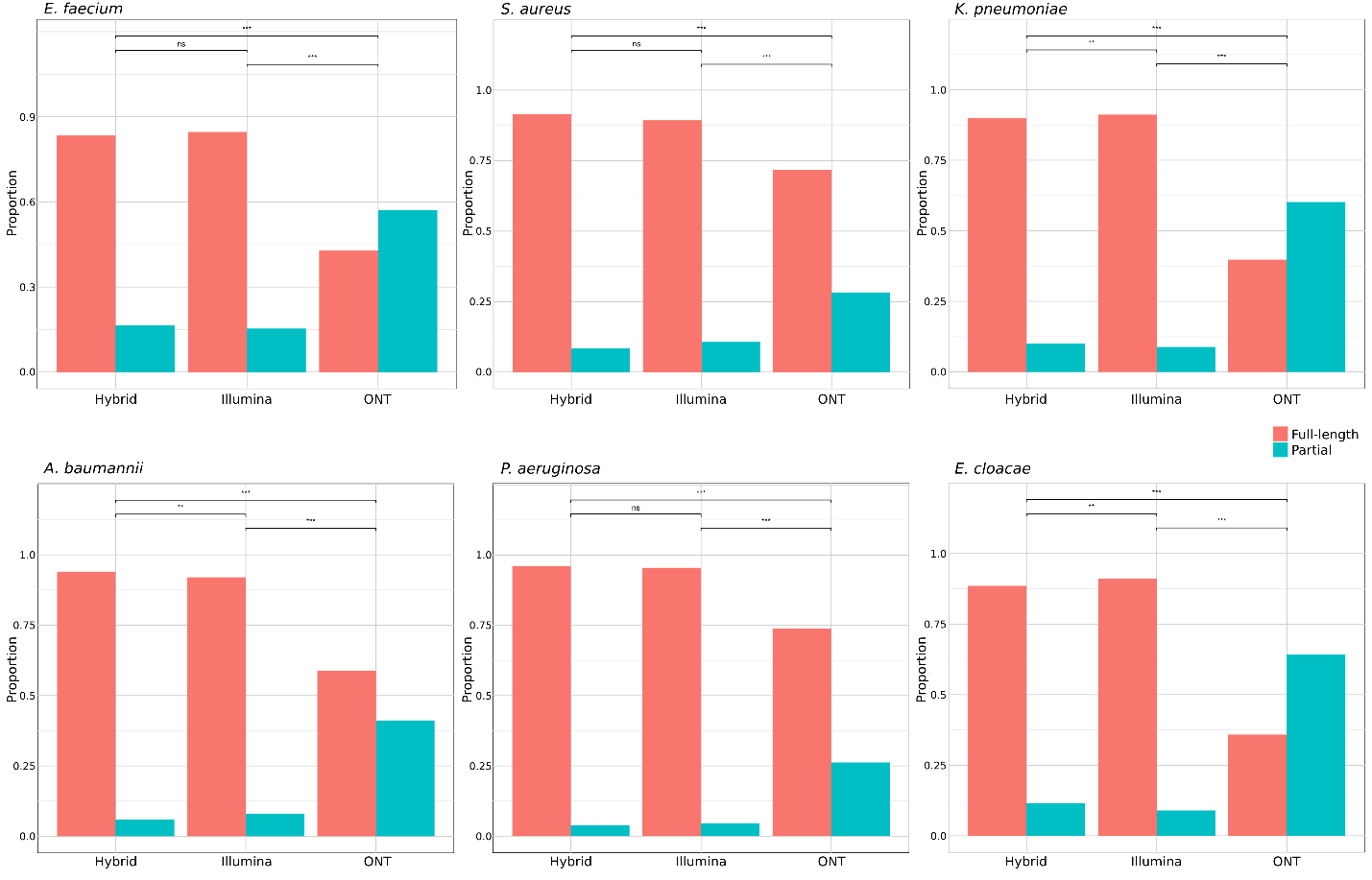

**Supplementary Figure 1**: Visualization of the differences in full-length (pink) versus partial genome (blue) hits for hybrid correction, Illumina and ONT across ESKAPE pathogens.

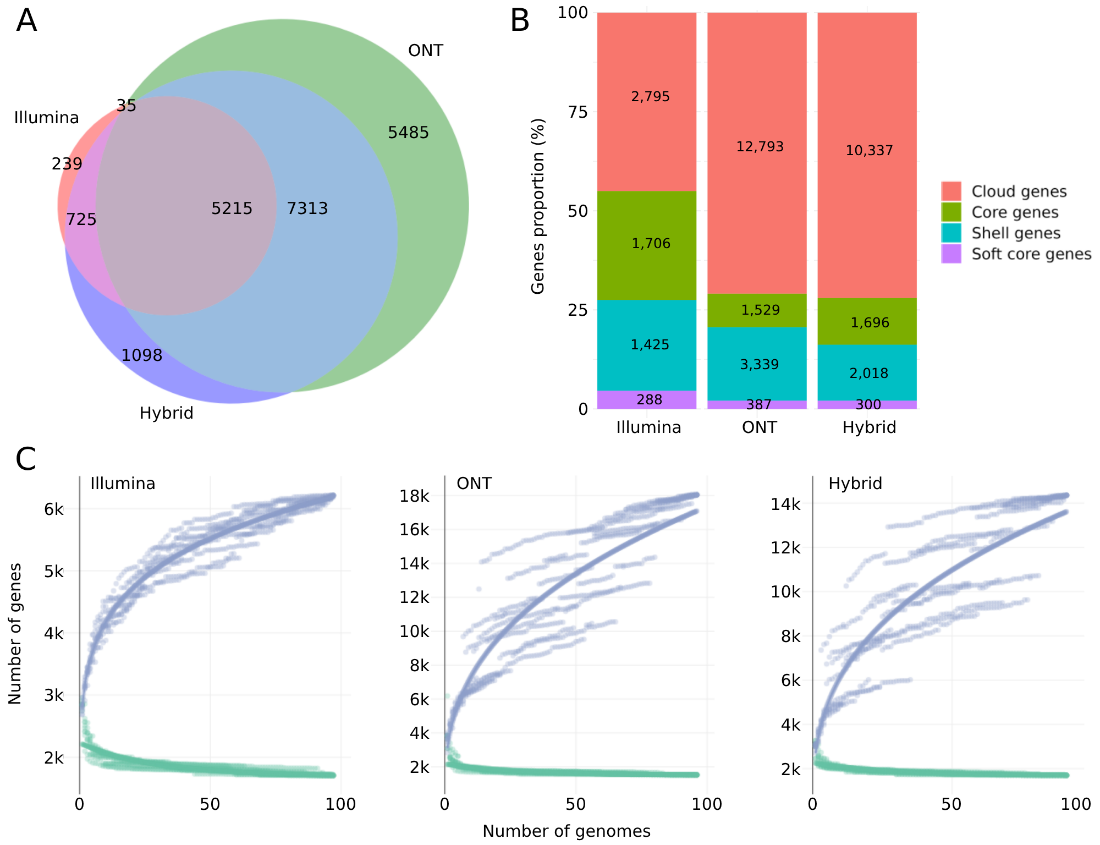

**Supplementary Figure 2**: Pangenome summary statistics for *E. faecium*. A represents the number of genes detected by Illumina, ONT and Hybrid correction both individually and overlapping between two or the three approaches. B represents the proportion of genes that characterize the core and accessory genes (shell, soft, and cloud), with the number within the bars representing the total number of genes within each section, while C shows the rarefaction curves of the relationship between the pangenome (blue) versus core genome (green).

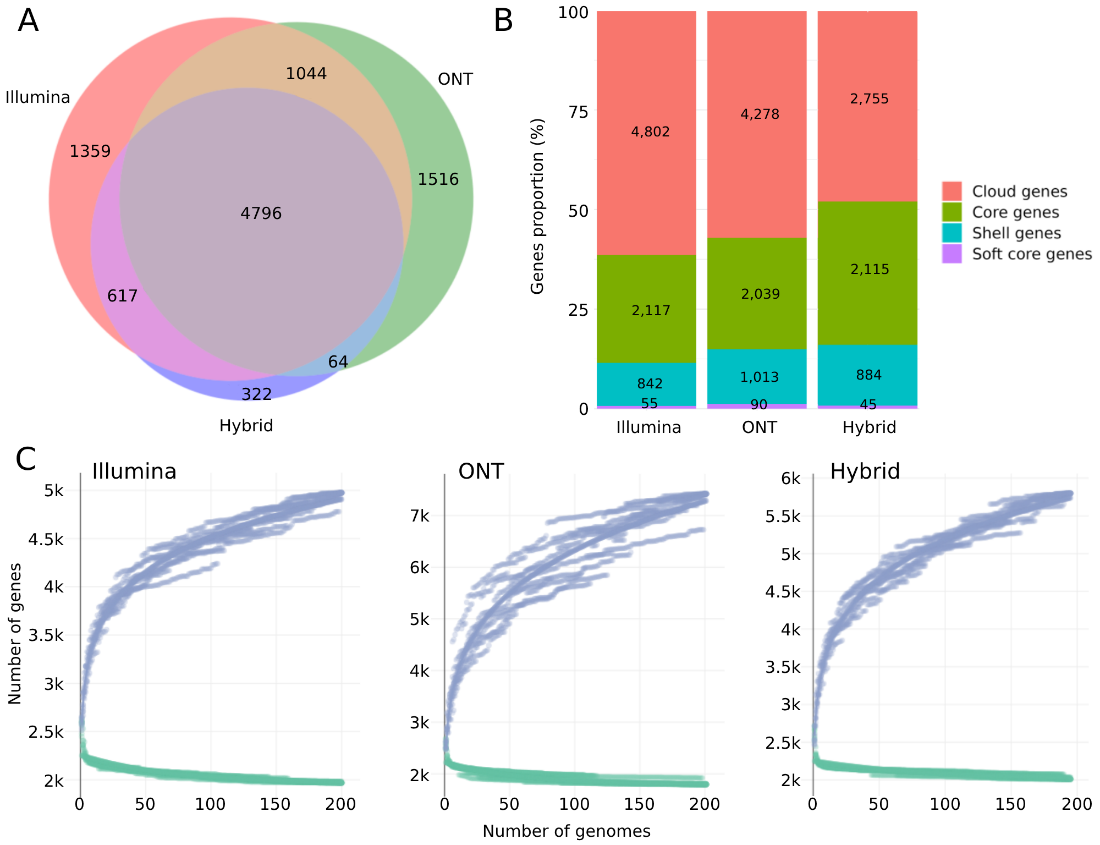

**Supplementary Figure 3:** Pangenome summary statistics for *S. aureus*. A represents the number of genes detected by Illumina, ONT and Hybrid correction both individually and overlapping between two or the three approaches. B represents the proportion of genes that characterize the core and accessory genes (shell, soft, and cloud), with the number within the bars representing the total number of genes within each section, while C shows the rarefaction curves of the relationship between the pangenome (blue) versus core genome (green).

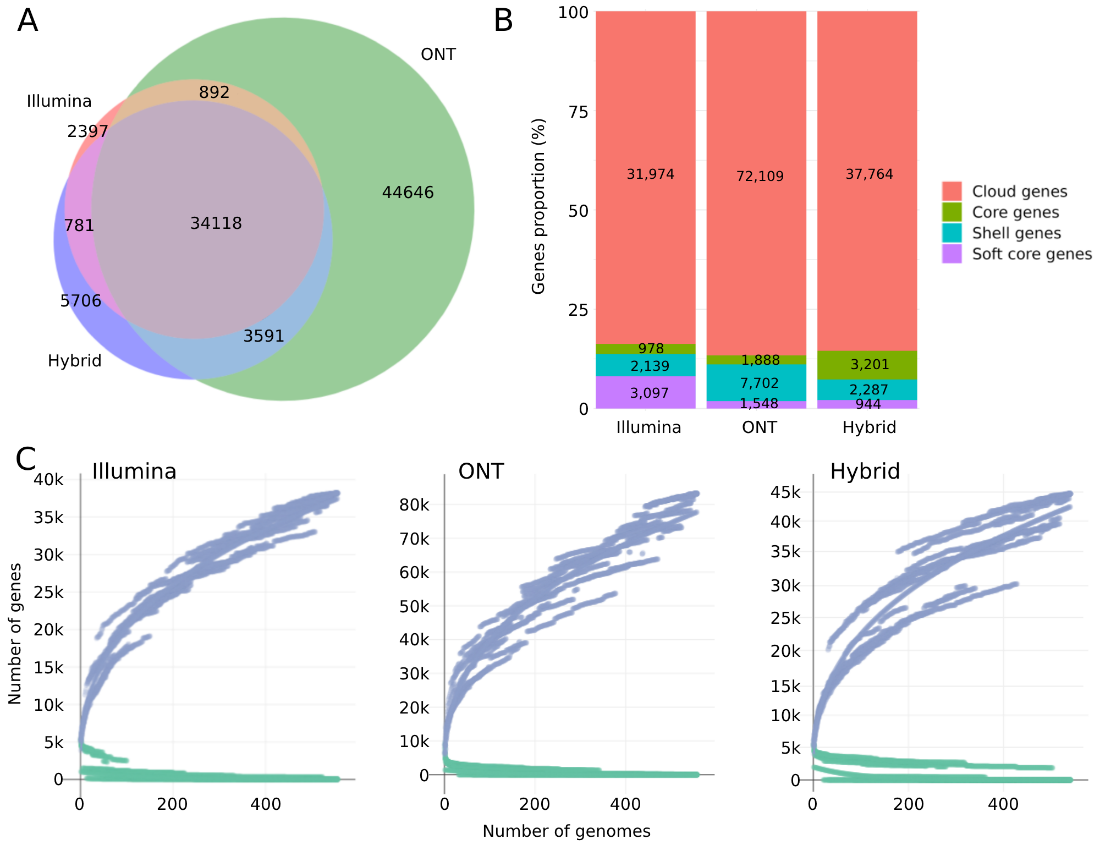

**Supplementary Figure 4:** Pangenome summary statistics for *K. pneumoniae*. A represents the number of genes detected by Illumina, ONT and Hybrid correction both individually and overlapping between two or the three approaches. B represents the proportion of genes that characterize the core and accessory genes (shell, soft, and cloud), with the number within the bars representing the total number of genes within each section, while C shows the rarefaction curves of the relationship between the pangenome (blue) versus core genome (green).

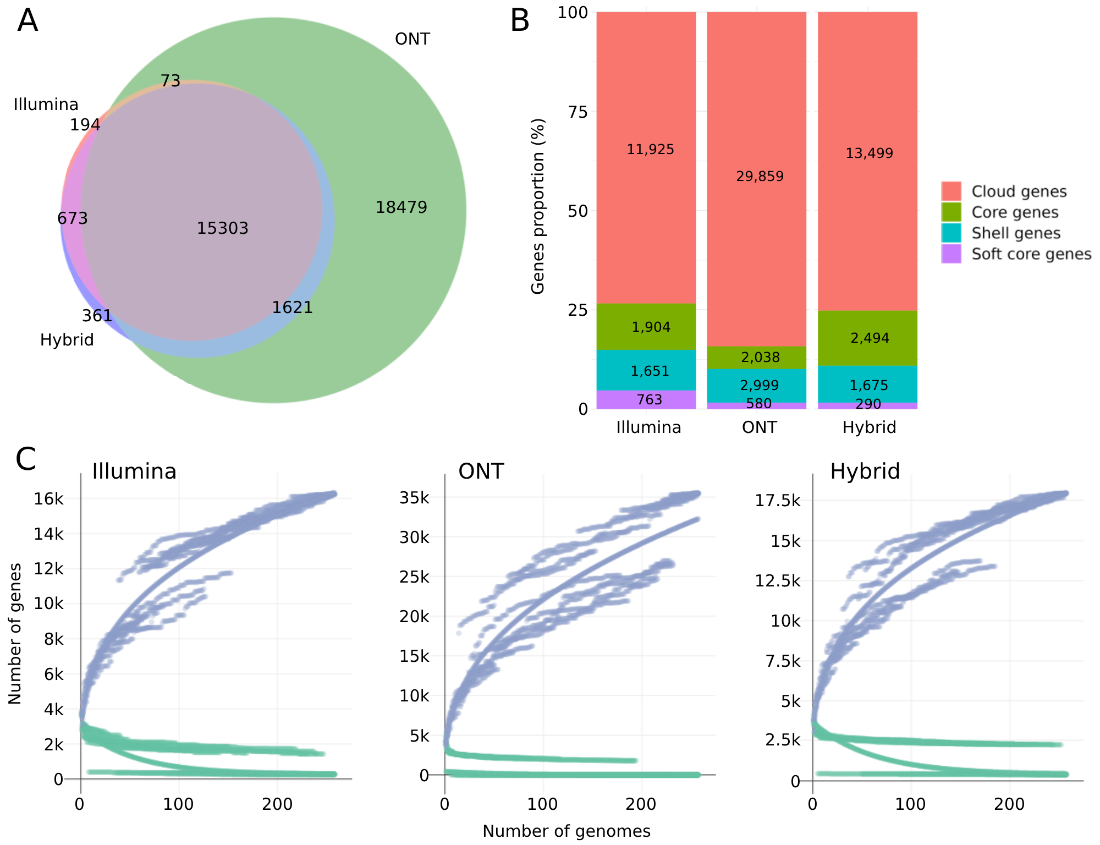

**Supplementary Figure 5:** Pangenome summary statistics for *A. baumannii*. A represents the number of genes detected by Illumina, ONT and Hybrid correction both individually and overlapping between two or the three approaches. B represents the proportion of genes that characterize the core and accessory genes (shell, soft, and cloud), with the number within the bars representing the total number of genes within each section, while C shows the rarefaction curves of the relationship between the pangenome (blue) versus core genome (green).

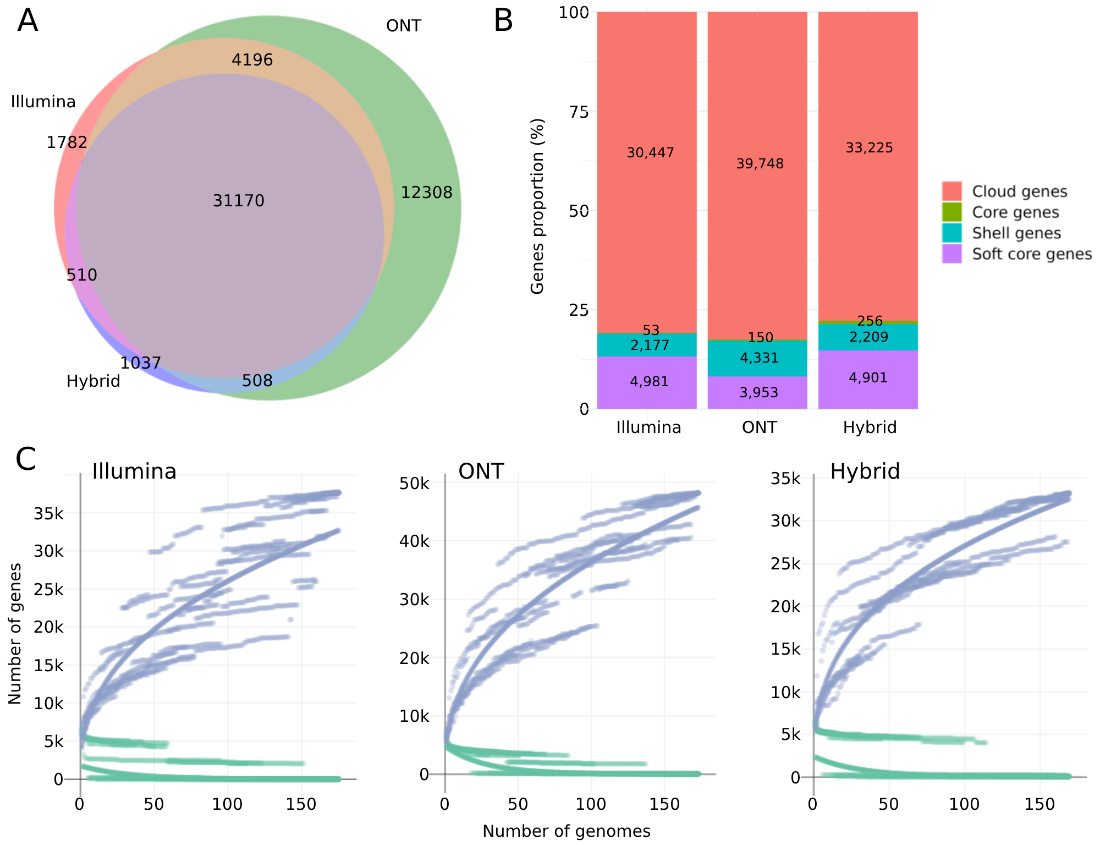

**Supplementary Figure 6:** Pangenome summary statistics for *P. aeruginosa*. A represents the number of genes detected by Illumina, ONT and Hybrid correction both individually and overlapping between two or the three approaches. B represents the proportion of genes that characterize the core and accessory genes (shell, soft, and cloud), with the number within the bars representing the total number of genes within each section, while C shows the rarefaction curves of the relationship between the pangenome (blue) versus core genome (green).

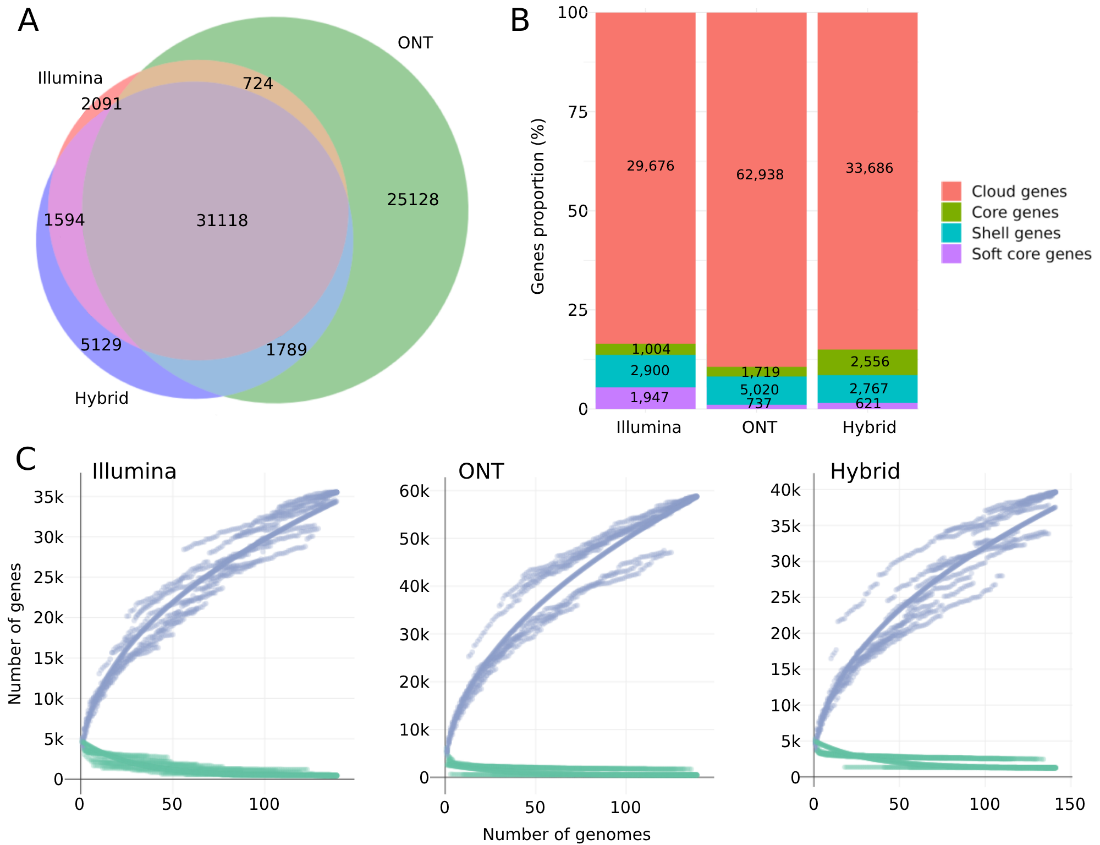

**Supplementary Figure 7**: Pangenome summary statistics for *E. cloacae*. A represents the number of genes detected by Illumina, ONT and Hybrid correction both individually and overlapping between two or the three approaches. B represents the proportion of genes that characterize the core and accessory genes (shell, soft, and cloud), with the number within the bars representing the total number of genes within each section, while C shows the rarefaction curves of the relationship between the pangenome (blue) versus core genome (green).

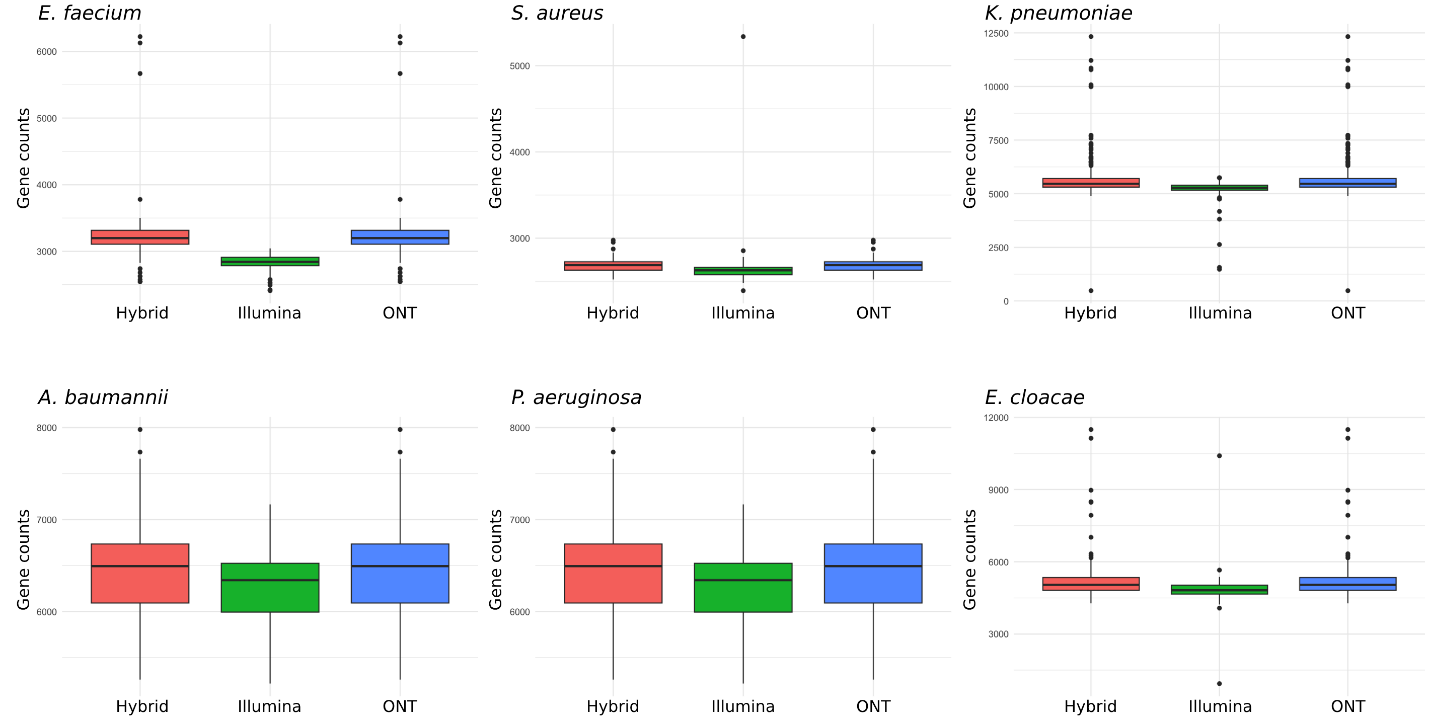

**Supplementary Figure 8**: Visualization of the comparison of the number of annotated genes prior to pangenome reconstruction detected by hybrid correction (pink), Illumina (green) and ONT (blue) across ESKAPE bacteria.
